# Opposing actomyosin pools generate cortical flows that establish epithelial polarity

**DOI:** 10.1101/2024.08.26.609703

**Authors:** Yu Shi, Lauren S. Ohler, Bailey N. de Jesus, Daniel J. Dickinson

## Abstract

Epithelial cell polarity, defined by distinct apical and basolateral domains, is fundamental for animal embryonic development and organ function. During organogenesis, epithelia often develop from unpolarized precursor cells. How mammalian epithelial cells establish polarity *de novo* from an initially unpolarized state has remained unclear, in part due to an inability to observe this process in real time in non-transformed cellular systems. Here, we leverage recent advances in 3D spheroid culture of mouse embryonic stem cells, fluorescent protein knock-in and live imaging techniques to study the process of epithelial polarity establishment. We show that apical myosin activity, regulated by Myosin Light Chain Kinase (MLCK), is crucial for the establishment and maintenance of epithelial polarity. Actomyosin cortical flows transport ZO-1, a tight junction component that interacts with apical polarity proteins, to establish the apical membrane. A second pool of myosin, regulated by Rho kinase, localizes basally and balances apically directed flow. Our result imply that epithelial polarity emerges as a consequence of actomyosin-driven cytoskeletal rearrangements.

## Introduction

Cell polarity, characterized by the asymmetric distribution of cellular components, plays a pivotal role in animal physiology (Campanale et al., 2017). Polarity not only determines cellular orientation but also drives crucial processes, from directed movement and morphogenesis to signal transduction and developmental differentiation. Epithelial cells, which make up most animal organs, establish distinct apical and basolateral membrane domains on opposite sides of the cell. Cells within an epithelium align their apico-basal axes, forming sheets of cells that can fold into spherical or tubular structures, foundational units of organs or tissues(Bryant and Mostov, 2008; Rodriguez-Boulan and Macara, 2014).

This epithelial organization, in which the apical membrane faces the external environment or lumen of an organ and the basolateral side contacts underlying extracellular matrix, is pivotal for organ function (Roignot et al., 2013). For instance, in the lungs, the apical surface of epithelial cells acts as a barrier against microorganisms, while the basolateral membrane helps in absorbing nutrients and hormones from the bloodstream (Kazmierczak et al., 2001; Sharma et al., 2020). In the intestines, the apical areas assist in nutrient uptake and set sites for digestive enzyme release, whereas the basolateral membrane aids in distributing nutrients to the bloodstream (Hooton et al., 2015; Co et al., 2019; Röder et al., 2014; Koepsell, 2020; Klunder et al., 2017). Similarly, in the kidneys, the apical side manages nutrient reabsorption, directing it to basal blood vessels (Nelson, 1993; Schlüter and Margolis, 2012).

Beyond its role in mature tissues, epithelial polarity plays a key role in morphogenesis during embryonic development. In both human and mouse development, the epiblast, which eventually develops into the fetus, comprises a disorganized cluster of cells in the pre-implantation stage. Epithelial polarity is established around the time of implantation, when the epiblast transforms into a single-layer epithelium (Bedzhov and Zernicka-Goetz, 2014; Shahbazi et al., 2017; Carleton et al., 2022; Wang et al., 2021). This polarized structure results in the formation of a lumen or pro-amniotic cavity apically and a connection to the extraembryonic tissue basally (Figure 1A, top).

**Figure 1:**
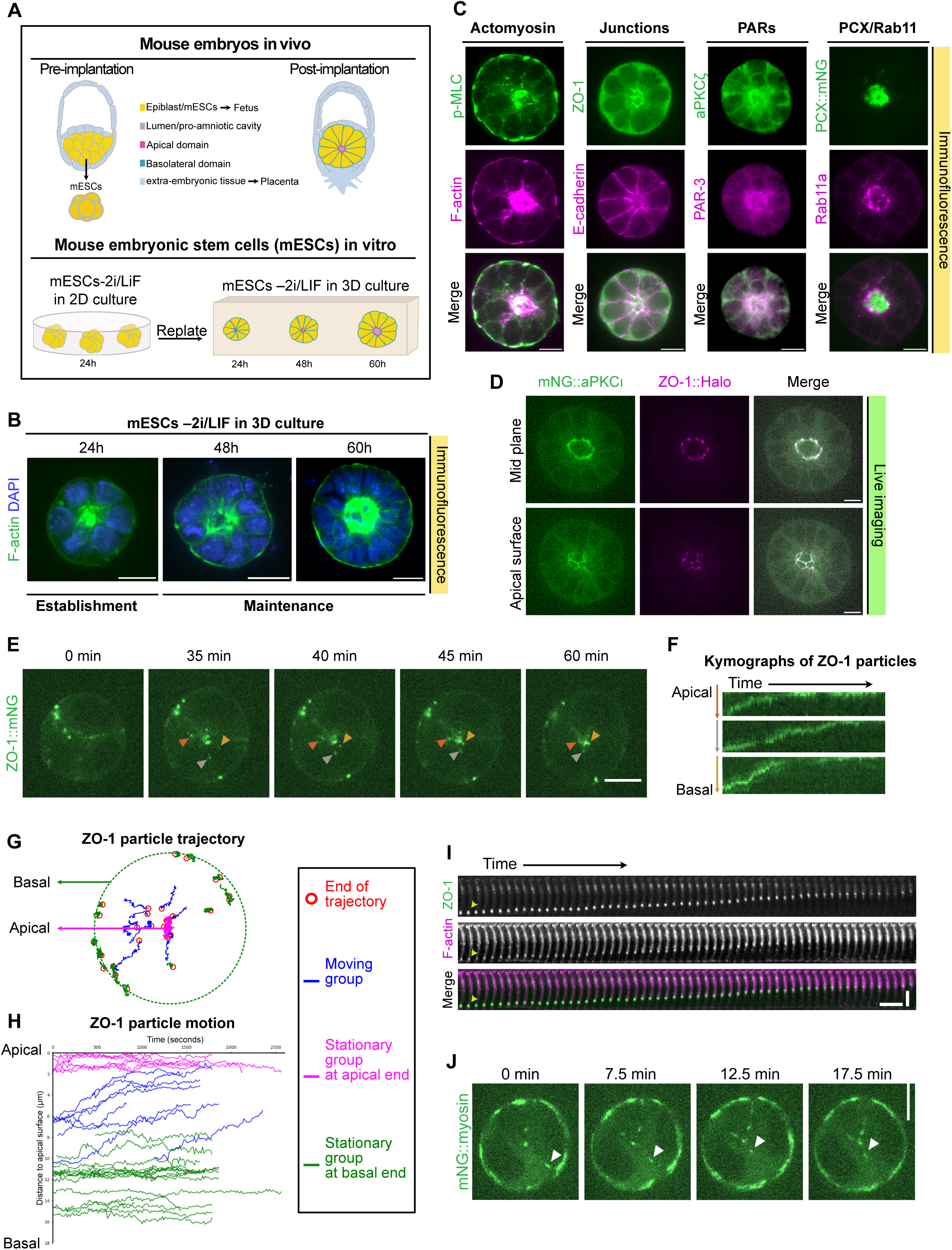
ZO-1 particles move apically during the establishment of epithelial polarity. (A) Illustration of epithelial polarity in mouse embryos *in vivo* (top) and the workflow of establishing epithelial polarity in mouse embryonic stem cells *in vitro* (bottom) (Bedzhov and Zernicka-Goetz, 2014). (B) Mid-plane images of fixed mESC spheroids in 3D culture at the indicated times after plating. Scale bars represent 10 μm. (C) Mid-plane images of fixed mESC spheroids stained with the indicated antibodies 48 hours after plating. Scale bars represent 10 μm. (D) Confocal images of mNG::aPKCι and ZO-1::Halo at two z planes (mid and apical surface plane) in dual-tagged mESCs 50 hours after plating. Scale bars represent 10 μm. (E) Confocal timelapse images of ZO-1 particles moving towards the future apical surface during polarity establishment. Arrowheads indicate particles whose movement was visualized in the kymographs in (F). Each image is a maximum-intensity projection of 3 planes spanning a total of 3 µm. Scale bars represent 10 μm. (F) Kymographs showing the movement of ZO-1 particles indicated in (E). The horizontal axis represents time, and each vertical axis shows the radial distance from the apical to the basal side of a cell. Colors match the arrowheads in (E). (G) Trajectories of 32 ZO-1 particles from 4 different spheres during polarity establishment. The observation times ranged from 27.5 to 42.75 minutes. There are 3 groups of trajectories: 7/32 stationary trajectories at the apical end (magenta), 17/32 stationary trajectories at the basal end (green), and 8/32 moving trajectories, which migrated from basal to apical (blue). Red circles indicate the ends of each trajectory. The basal surface is represented by a green dash circle (normalized radius = 1) and the apical surface is at the center. (H) Plot of position as a function of time for each tracked ZO-1 particle represented in (G). (I) Montage showing the concordant movement of ZO-1 and F-actin toward the apical surface in a polarizing mESC spheroid. Vertical scale represents 5 µm, and horizontal scale bar represents 90s. (J) Confocal live images showing the movement of myosin during polarity establishment. The white arrowhead indicates a moving myosin particle. Each image is a maximum-intensity projection of 5 planes spanning a total of 6 µm. Scale bars represent 10 μm.

Widely studied epithelial models, such as MDCK cells, have revealed that the establishment of epithelial polarity necessitates a coordinated interplay of cytoskeleton, membrane trafficking, adhesion and cellular signaling (Martin et al., 2021; Rodriguez-Boulan and Macara, 2014). For example, the actomyosin network, encompassing actin filaments (F-actin) and non-muscle myosin II, collaboratively shape cells (Raman et al., 2018; Miao and Blankenship, 2020). Junction proteins like the tight junction marker ZO-1 and adherens junction marker E-cadherin facilitate intercellular connections (Niessen, 2007). Signaling components such as the partitioning defective proteins or PARs, including the kinase aPKC and the scaffold protein PAR-3, serve as polarity determinants (Horikoshi et al., 2009). The apical anti-adhesive sialomucin Podacalyxin (PCX) plays a key role in lumen formation (Desclozeaux et al., 2008; Meder et al., 2005; Bryant et al., 2010). Additionally, the orientation of epithelial polarity depends on interactions between cells and extracellular matrix (ECM)(Yu et al., 2005; Bryant et al., 2014).

Recent work in mouse embryos and embryonic stem cells have underscored the ECM’s role in providing spatial cues for polarization (Bedzhov and Zernicka-Goetz, 2014) and the importance of cadherin-mediated cell adhesion for establishing the initial polarity axis (Liang et al., 2022). While recent advances have expanded our knowledge, we still lack a comprehensive understanding of the mechanisms underlying *de novo* epithelial polarity establishment. This gap in knowledge is primarily due to the lack of dynamic data, with limitations in knock-in reporter systems and time-lapse imaging techniques posing significant challenges.

To unravel the process of epithelial polarity establishment, it is crucial to understand how polarity proteins are transported to their designated polarized domains. Non-muscle myosin II (hereafter referred to as myosin) generates local contractile forces on F-actin, which can lead to large-scale cortical flows that transport cortical proteins within cells. In one widely-studied invertebrate polarity model, the *C. elegans* zygote, myosin activity generates cortical flows that transport PARs to establish the polarity axis (Goehring et al., 2011; Munro et al., 2004; Mittasch et al., 2018; Chang and Dickinson, 2022; Dickinson et al., 2017; Motegi and Sugimoto, 2006; Cheeks et al., 2004; Motegi et al., 2011; Illukkumbura et al., 2023; Mayer et al., 2010). There is evidence that similar myosin-driven cortical flows contribute to polarity establishment in *Drosophila* neuroblasts (LaFoya and Prehoda, 2021). In mammals, existing knowledge indicates that a polarized actomyosin network is essential for forming apical PAR protein caps in 8-cell mouse embryos (Zhu et al., 2017). Additionally, myosin, along with components of adherens and tight junctions, is enriched at cell contacts to facilitate the sealing of the embryo, a critical step for mouse blastocyst formation (Zenker et al., 2018). However, whether myosin activity plays a role in transporting junction components or PARs in vertebrate polarity models, particularly in epithelial polarity, is unclear.

To probe this question, we utilized a 3D spheroid culture of mouse embryonic stem cells (mESCs) as our model to study epithelial polarity. These cultures mimic the epithelial polarization seen in early-stage mouse embryos, providing an opportunity to study how an organized epithelial tissue emerges from previously unpolarized cells. We have developed an effective and rapid fluorescent protein knock-in technique for mESCs (Shi et al., 2023), enabling us to monitor protein dynamics using live imaging without overexpression artifacts. Here we show that during epithelial polarity establishment in mESCs, junctional proteins, most notably ZO-1, are carried towards the apical region by directional actomyosin flows. Using pharmacological inhibitors, we show that apical actomyosin flows require Myosin Light Chain Kinase (MLCK). These flows, in turn, and are essential for polarization of tight junctions, PARs, and the luminal component PCX. A second, Rho kinase (ROCK)-dependent pool of myosin is active at the basal surface, and ROCK activity is necessary for exclusion of apical proteins from the basal domain. Together, these results reveal roles for opposing actomyosin pools in epithelial cells and shed light on the process of *de novo* epithelial cell polarization in mammalian embryos.

## Results

### ZO-1 particles flow towards the apical surface during polarity establishment

We utilized 3D spheroid cultures of mESCs to explore epithelial polarity. mESCs come from pre-implantation mouse epiblasts and are cultured under conditions that maintain naïve pluripotency (2i/LIF) (Mulas et al., 2019). Upon withdrawal of 2i/LIF, mESCs exit naïve pluripotency and enter a state termed formative pluripotency (Hayashi et al., 2011; Kalkan et al., 2017; Mulas et al., 2017; Kinoshita et al., 2021), in which they can undergo epithelial polarization similar to mouse epiblasts around the time of implantation *in vivo* (Bedzhov and Zernicka-Goetz, 2014; Shahbazi et al., 2017) (Figure 1A, top). We cultured mESCs in a 2D environment for 24 hours following withdrawal of 2i/LIF (Figure 1A, bottom), then transferred the cells to 3D culture conditions using Matrigel, which provides extracellular matrix cues that support epithelial polarization (Bedzhov and Zernicka-Goetz, 2014). We allowed cells to grow in the 3D setting for up to 60 hours, during which they gradually developed into a polarized spheroid with an apical lumen in the center (Figure 1A, bottom & 1B). Note that in previous reports, cells grown under similar conditions have been described as having formative, primed or epiblast-like pluripotent characteristics. We did not attempt to distinguish these different states here; for simplicity, we refer to cells cultured for >24h in the absence of 2i/LIF as “primed mESCs.”

To explore epithelial polarity establishment in mESCs and confirm previous findings (Shahbazi et al., 2017; Bedzhov and Zernicka-Goetz, 2014; Shi et al., 2023), we fixed 3D cultures after polarity establishment and performed immunofluorescence staining. As expected, we observed apical enrichment of several proteins, including phosphorylated myosin light chain (p-MLC), F-actin, ZO-1, aPKCζ, PAR-3, PCX, and Rab11a (Figure 1C). E-cadherin was found at cell-cell contacts. Notably, we observed that p-MLC accumulated apically, but also in patches at the basal cell surface, consistent with our previous report (Figure 1C)(Shi et al., 2023). Next, to visualize polarity establishment in live cells, we used a streamlined CRISPR knock-in protocol (Shi et al., 2023) to endogenously tag the key polarity kinase aPKC (mNeonGreen::aPKCι or mNG::aPKCι; Figure S1A). In the same cells, we also tagged the tight junction protein ZO-1 (ZO-1::Halo). In live cells, we observed a striking colocalization of aPKC and ZO-1 at the margins of the apical domain (Figure 1D, Movie 1), consistent with reports in immortalized epithelial cell lines (Ikenouchi et al., 2007; Hurd et al., 2003; Whyte et al., 2010; Mangeol et al., 2022). These data suggest a potential association of ZO-1 with aPKC at the apical membrane in polarizing mESCs. These findings confirm coordinated translocation of cytoskeletal, membrane and polarity signaling proteins to the apical domain in primed mESCs in 3D culture.

Next, we asked how apicobasally polarized protein localization is established. We performed live imaging of primed mESCs as they transitioned from a disorganized to a polarized state. Polarity establishment occurred at different times in different cell clusters, but typically lasted 2-4 hours and happened 24–48 hours after plating in 3D culture. Strikingly, we observed that during polarity establishment, endogenous ZO-1 formed clusters that traveled along cell-cell contacts towards the apical membrane (Figure 1E & 1F, Movie 2). To quantify this behavior, we tracked single ZO-1 particles over time (Jaqaman et al., 2008; Ershov et al., 2022). Across 4 polarizing spheroids, 25% of ZO-1 clusters were moving towards the apical membrane, while the remainder were stationary at either the apical or basal surface (Figure 1G-H & S1B-C). Since ZO-1 is an actin binding protein (Belardi et al., 2020; Fanning et al., 1998; Odenwald et al., 2016), we hypothesized that cortical F-actin flows might be responsible for ZO-1 movement. Indeed, when we imaged ZO-1 and F-actin together by labeling ZO-1::mNG knock-in cells with a fluorogenic live-cell F-actin dye (Lukinavičius et al., 2014), we saw that moving ZO-1 particles always colocalized with moving F-actin punctae (9/9 polarizing spheroids; Figure 1I and Movie 3). We also observed apically directed movements of non-muscle Myosin II (Figure 1J) and the adherens junction marker β-catenin (Figure S1D). Together, these data suggested the presence of an apically directed cortical flow that transports junctional proteins during polarity establishment.

### MLCK-driven myosin activity is necessary for forming the apical ZO-1 domain

Cortical flows, powered by myosin contractility, are instrumental in transporting polarity proteins to establish asymmetry in invertebrate model systems (Munro et al., 2004; Motegi and Sugimoto, 2006; LaFoya and Prehoda, 2021). Based on this, we hypothesized that in mESCs, myosin activity could lead to an apical-directed flow that might drive ZO-1 particles towards the apical region, thereby facilitating epithelial polarity establishment. To test this hypothesis, we treated cells with ML-7, a potent and specific inhibitor that targets Myosin Light Chain Kinase (MLCK), thereby blocking myosin phosphorylation and activation (Martinsen et al., 2014; Xiong et al., 2017). We observed cells within the first hour of ML-7 treatment to identify immediate effects, avoiding longer treatments due to reported apoptosis caused by extended ML-7 exposure (Connell and Helfman, 2006).

Treatment with ML-7 strikingly disrupted the apical accumulation of ZO-1 in polarizing cells. ZO-1 particles started retracting from the apical region within 10 minutes post ML-7 treatment, with further retraction and cell shape distortion observed from 30 to 60 minutes (Figure 2A-B, Movie 4), while the ZO-1 apical domain remained intact following DMSO treatment (Figure S2A). Nuclei maintained a normal appearance following ML-7 treatment (Figure S2B), suggesting that loss of polarity was not due to apoptosis. Using particle tracking, we found that the majority of tracked particles (73%) moved towards the basal surface following ML-7 treatment (Figure 2C-D, Figure S2C-D). A smaller fraction of clusters moved in the apical direction (8%), while some remained stationary (19%) following ML-7 treatment (Figure 2D & S2D). The disruption of apical ZO-1 transport by ML-7 was reversible: apical ZO-1 recovered following washout (Figure 2E-G, Movie 5). To quantitatively assess this disruption and recovery of apical ZO-1 in response to ML-7 and wash-off treatment, we measured the apical ZO-1 intensity from mid-plane confocal sections (Figure S2E). The fraction of mNG::ZO-1 signal found in the apical region did not change following DMSO treatment (Figure 2E-F, Figure S2F, top). In contrast, treatment with ML-7 led to a significant decrease in apical ZO-1, while its levels significantly increased following wash-off (Figure 2E-F, Figure S2F, bottom). Together, these data show that myosin activity downstream of MLCK is essential for the directional movement of ZO-1 particles and for the establishment and maintenance of the apical ZO-1 domain.

**Figure 2:**
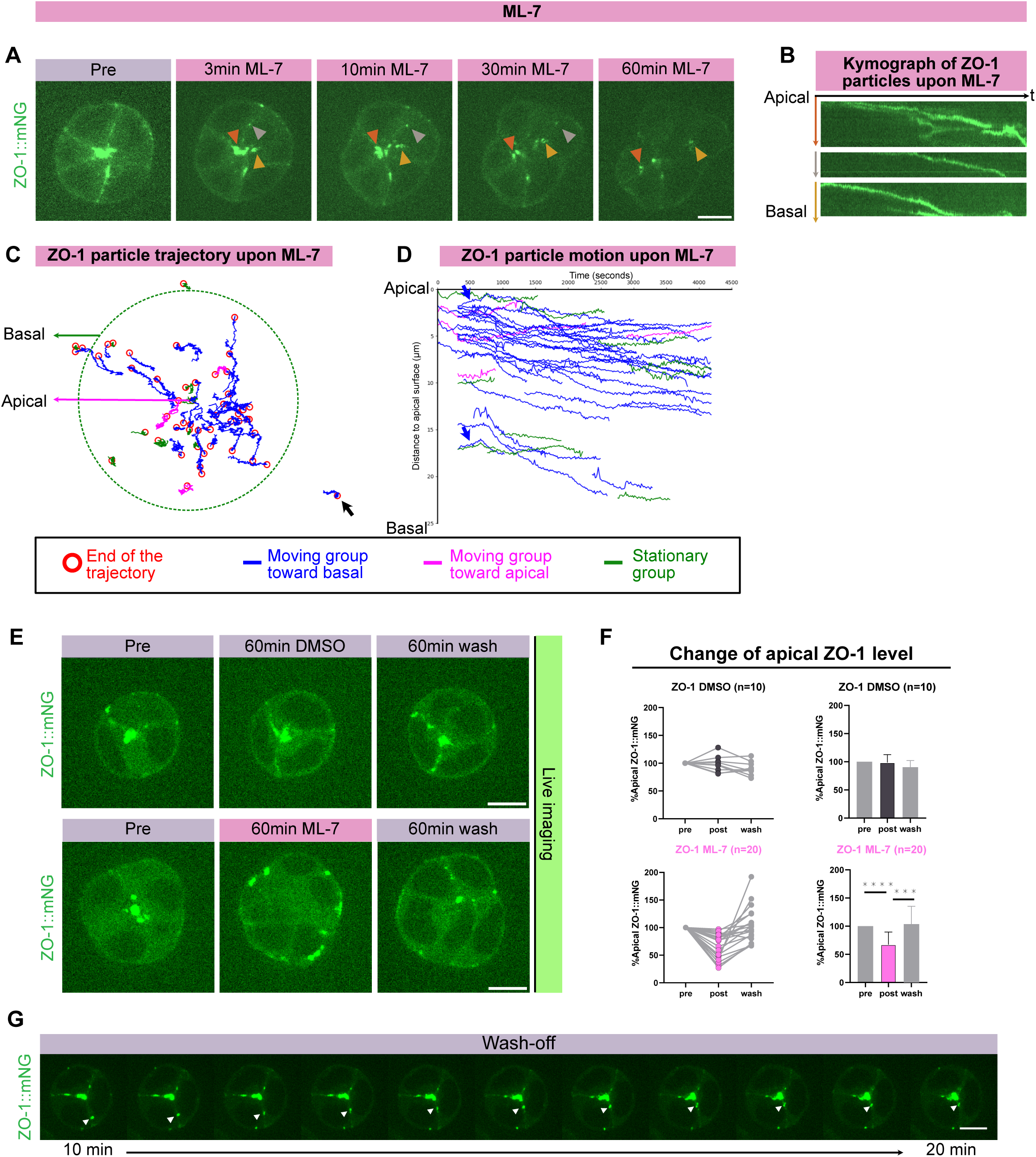
The formation of the apical ZO-1 domain depends on MLCK-driven myosin activity. (A) Confocal timelapse images of ZO-1 particles moving towards basal surface upon ML-7 treatment during polarity establishment. Arrowheads indicate particles whose movement was visualized in the kymographs in (B). Each image is a maximum-intensity projection of 3 planes spanning a total of 3 µm. (B) Kymographs showing the movement of these ZO-1 particles from (A). The horizontal axis represents time, and each vertical axis shows the radial distance from the apical to the basal side of a cell. Colors match the arrowheads in (A). (C) Trajectories of 52 ZO-1 particles from 3 different spheres during polarity establishment. The observation times were 70 minutes. There are 3 groups of trajectories: 38/52 was basal-moving group which migrated from apical to basal towards (blue), 4/52 was apical-moving group (magenta) and 10/52 was stationary group (green). The red circles indicate the ends of each trajectory. The basal surface is represented by a green dash circle (normalized radius = 1) and the apical surface is at the center. The black arrow represents one basal-moving trajectory outside the original basal surface due to a distorted shape (radius > 1). (D) Plot of position as a function of time for each tracked ZO-1 particle represented in (C). Two blue arrows depict trajectories moving from basal to apical, then retracting back to basal. (E) Mid-plane images of live ZO-1::mNG spheroids in 3D culture during polarity establishment. Top panel displays three stages of DMSO treatment: pre-treatment, 60-minute post-treatment, and 60 minutes after wash-off. Bottom panel shows three stages of ML-7 treatment: pre-treatment, 60-minute post-treatment, and 60 minutes after wash-off. (F) Normalized percentage of apical ZO-1::mNG at the midplane assessed before treatment, 60 minutes after ML-7 treatment, and 60 minutes following the wash-off, for groups treated with DMSO (n = 10) and ML-7 (n = 20). Data are normalized to corresponding pre-treatment levels; for non-normalized data, see Figure S2F. Left panel displays data for individual observations at three stages; right panel shows the mean data with standard deviation. Asterisks indicate significant changes (paired t-test; see Table S1 for exact p values). (G) Confocal timelapse images of ZO-1 particles from 10 to 20 minutes upon wash-off following 60-minute ML-7 treatment. Arrowheads indicate a particle’s apical movement. Time lapse are at 1-minute interval. Each image is a maximum-intensity projection of 3 planes spanning a total of 3 µm. Scale bars represent 10 μm.

### MLCK-driven myosin activity is required for apical membrane formation

We next asked if MLCK-regulated myosin activity also influences other apical polarity proteins, including PARs and trafficking components, or alternatively if the effect of myosin is specific to ZO-1. First, using immunofluorescence, we assessed the polarization of PCX and aPKCζ alongside ZO-1 upon ML-7 treatment. As expected, both PCX and aPKCζ localized in the apical domain in controls (Figure 3A, top). However, after a 60-minute ML-7 treatment, the polarized distribution PCX and aPKCζ in the apical domain was disrupted along with ZO-1 (Figure 3A, bottom). DAPI staining confirmed intact nuclear shapes, ruling out ML-7-induced apoptosis as the cause for this disruption (Figure 3A). This suggests that MLCK-regulated myosin activity is essential for the apical distribution of PARs and luminal proteins in addition to ZO-1.

**Figure 3:**
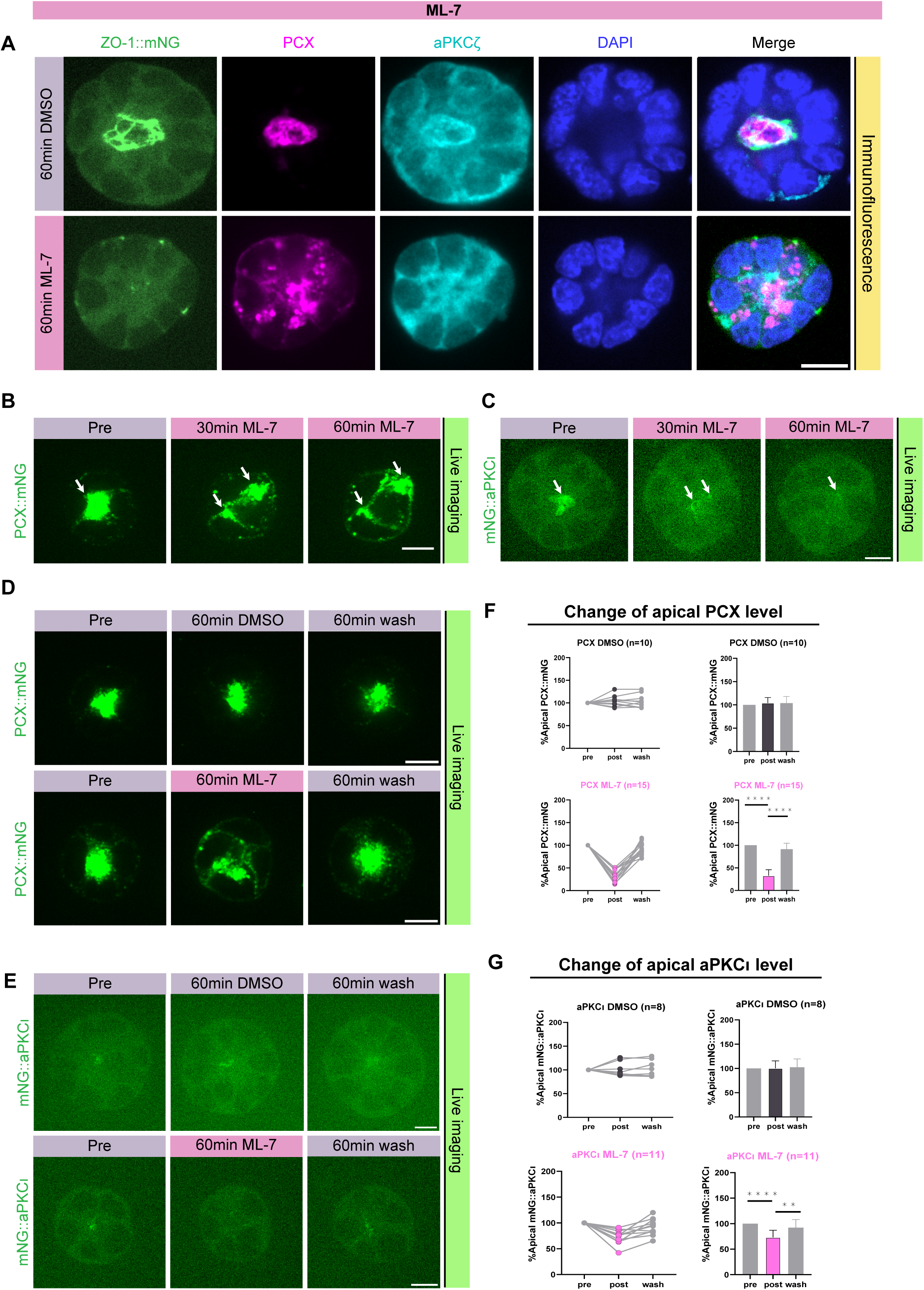
MLCK-driven myosin activity is essential for the apical domain formation of both aPKC and PCX. (A) Mid-plane images of fixed ZO-1::mNG spheroids with the indicated antibodies 48 hours after plating. The top panel shows spheroids treated with 60 minutes of DMSO, and the bottom panel with 60 minutes of ML-7. (B) Confocal timelapse images of PCX particles moving towards basal surface upon ML-7 treatment during polarity establishment. Arrows indicate the split particles. Each image is a maximum-intensity projection of 3 planes spanning a total of 3 µm. (C) Mid-plane images of live mNG::aPKCι spheroids treated with 60-minute ML-7 during polarity establishment. Arrows indicate the change of aPKCι domain. (D) Mid-plane images of live PCX::mNG spheroids in 3D culture during polarity establishment. Top panel displays three stages of DMSO treatment: pre-treatment, 60-minute post-treatment, and 60 minutes after wash-off. Bottom panel shows three stages of ML-7 treatment: pre-treatment, 60-minute post-treatment, and 60 minutes after wash-off. (E) Mid-plane images of live mNG::aPKCι spheroids in 3D culture during polarity establishment. Top panel displays three stages of DMSO treatment: pre-treatment, 60-minute post-treatment, and 60 minutes after wash-off. Bottom panel shows three stages of ML-7 treatment: pre-treatment, 60-minute post-treatment, and 60 minutes after wash-off. (F) Normalized apical PCX::mNG percentages at the midplane, measured pre-treatment, 60 minutes after treatment, and 60 minutes post-wash. The top panel relates to DMSO (n = 10) treatment, the bottom to ML-7 (n = 15), with all data adjusted to pre-treatment values; for non-normalized data, see Figure S3C. The left panel presents individual data points at the three stages; the right panel depicts average values with standard deviation. Significant changes are marked by asterisks (paired t-test; see Table S1 for exact p values). (G) Normalized apical mNG::aPKCι percentages at the midplane, measured pre-treatment, 60 minutes after treatment, and 60 minutes post-wash. The top panel relates to DMSO (n = 8) treatment, the bottom to ML-7 (n = 11), with all data adjusted to pre-treatment values; for non-normalized data, see Figure S3D. The left panel presents individual data points at the three stages; the right panel depicts average values with standard deviation. Significant changes are marked by asterisks (paired t-test; see Table S1 for exact p values). Scale bars represent 10 μm.

Building on our observations from fixed cells, we sought to dynamically track how these proteins respond to ML-7 treatment in live cells. We expressed PCX::mNG using an exogenous transgene and endogenously tagged mNG::aPKCι (Figure 1D). ML-7 treatment caused the single apical PCX domain to split and move towards the basal side of the cell (Figure 3B & S3A, Movie 6), and the apical enrichment of aPKCι diminished over time (Figure 3C & S3B). The nuclear shape remained consistent (Figure S3A & S3B). Wash-off experiments demonstrated that the loss of apical membrane identity following ML-7 treatment was reversible (Figure 3D & 3E, bottom). Quantitative assessment confirmed that ML-7 exposure led to a substantial decrease in apical PCX and aPKCι levels, which recovered to initial levels upon wash-off (Figure 3F & 3G, Figure S3C & S3D, bottom). Conversely, the apical distribution of PCX and aPKCι signal remained stable during DMSO treatment (Figure 3F & 3G, Figure S3C & S3D, top). These data show that MLCK-driven myosin activity is required for apical localization of PCX and aPKC.

### MLCK inhibition specifically targets the apical pool of myosin

Although the results of ML-7 treatment supported our hypothesis that myosin flows are important to establish apical ZO-1 localization and apicobasal polarity, we were surprised by the basal movement of ZO-1 that we observed following MLCK inhibition. Our prediction had been that ZO-1 would stop moving entirely if we blocked myosin activation; instead, we observed movement, but in the opposite direction from controls (Figure 2A-D). F-actin also redistributed to more basal and punctate localization following ML-7 treatment (Figure 4A & 4C, Movie 7). Interestingly, ZO-1 and F-actin were redistributed together, and still colocalized, following treatment with ML-7, suggesting that the basal redistribution of ZO-1 after MLCK inhibition is mediated by cytoskeletal changes.

**Figure 4:**
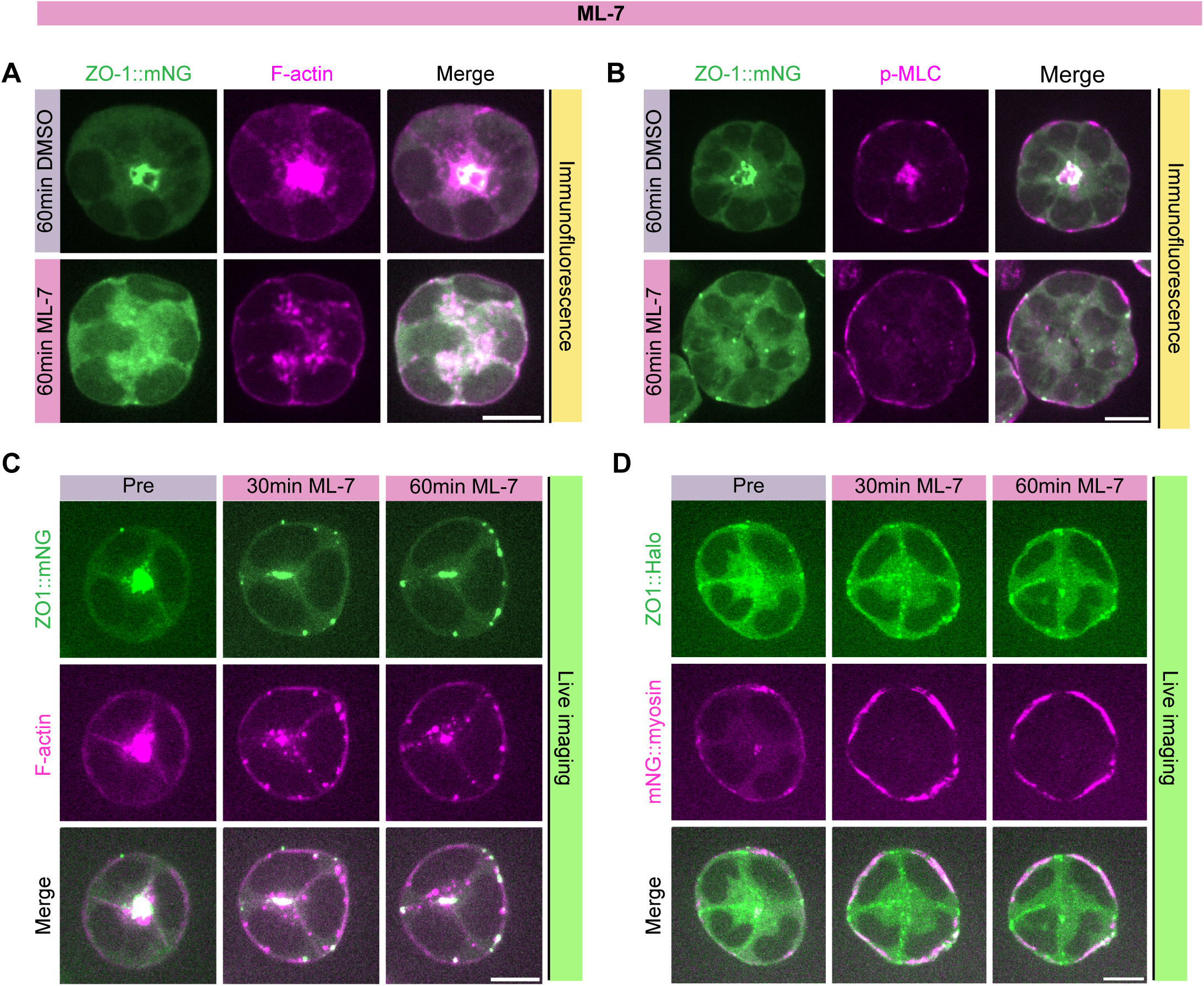
Inhibition of MLCK selectively affects apical myosin pool. (A) Mid-plane images of fixed ZO-1::mNG spheroids with Phalloidin 48 hours after plating. The top panel shows spheroids treated with 60 minutes of DMSO, and the bottom panel with 60 minutes of ML-7. (B) Mid-plane images of fixed ZO-1::mNG spheroids with the indicated antibody 48 hours after plating. The top panel shows spheroids treated with 60 minutes of DMSO, and the bottom panel with 60 minutes of ML-7. (C) Confocal timelapse images of ZO-1 and F-actin upon ML-7 treatment during polarity establishment. Each image is a maximum-intensity projection of 3 planes spanning a total of 3 µm. (D) Confocal timelapse images of ZO-1 and myosin upon ML-7 treatment during polarity establishment. Each image is a maximum-intensity projection of 3 planes spanning a total of 3 µm. Scale bars represent 10 μm.

Next, we examined the organization of myosin. We expected that MLCK inhibition would eliminate phospho-myosin (p-MLC) staining. Surprisingly, while apical p-MLC was abolished by ML-7 treatment, the basal p-MLC signal remained (Figure 4B). Using dual-tagged mESCs (Figure S4), live imaging of endogenously tagged myosin and ZO-1 (mNG::myosin and ZO-1::Halo) confirmed that the apical pool of cortical myosin, but not the basal pool, was specifically disrupted by MLCK inhibition (Figure 4D). These observations suggest that MLCK primarily regulates a unique apical actomyosin pool, while the basal pool is regulated by a different mechanism. The basal pool of actomyosin that remains after MLCK inhibition could potentially account for the basal flow of ZO-1 following ML-7 treatment.

### Inhibition of ROCK selectively affects the basal myosin pool

How is basal myosin activation sustained in the absence of MLCK activity? The Rho-associated protein kinase (ROCK), which is known to regulate actomyosin organization, was an attractive candidate regulator (Julian and Olson, 2014). ROCK can both phosphorylate MLC directly and inhibit myosin phosphatase, thereby enhancing myosin activity. Previous studies indicate that ROCK and MLCK regulate myosin activity in opposite spatial domains within fibroblasts (Totsukawa et al., 2000, 2004). Consequently, we hypothesized that ROCK might predominantly influence myosin activity at the basal surface of polarizing mESCs. To test this hypothesis, we employed H1152, a specific ROCK inhibitor (Sasaki et al., 2002). We avoided the more widely used compound Y27632 for these experiments because Y27632 can also inhibit aPKC (Atwood and Prehoda, 2009).

We tested the impact of H1152 on both myosin and ZO-1. Prior to treatment, basal myosin patches were clearly visible, and ZO-1 was predominantly located at the apical surface (Figure 5A, left). Upon introducing H1152, the basal myosin disappeared immediately, while ZO-1’s apical localization was progressively disrupted over time (Figure 5A, middle, Figure S5A-C). Once H1152 was removed, basal myosin quickly reappeared, and in tandem, the basal mislocalization of ZO-1 began to reduce (Figure 5A, right and Figure S5C). Overall, while myosin’s distribution underwent swift changes, ZO-1’s shifts were more gradual, in the presence of H1152 or after its removal. Immunofluorescence confirmed that ROCK inhibition specifically disrupted basal myosin activation, as visualized by p-MLC staining (Figure 5B). Notably, the apical localization of myosin and p-MLC remained unchanged despite of H1152 treatment (Figure 5A-B). To further support this, the quantification of apical mNG::myosin signal in live cells confirmed that its levels were not affected by H1152 treatment (Figure 5C, S4D, bottom). These findings suggest that ROCK specifically regulates a basal pool of active myosin, additionally, prevents ZO-1 from localizing at the basal membrane.

**Figure 5:**
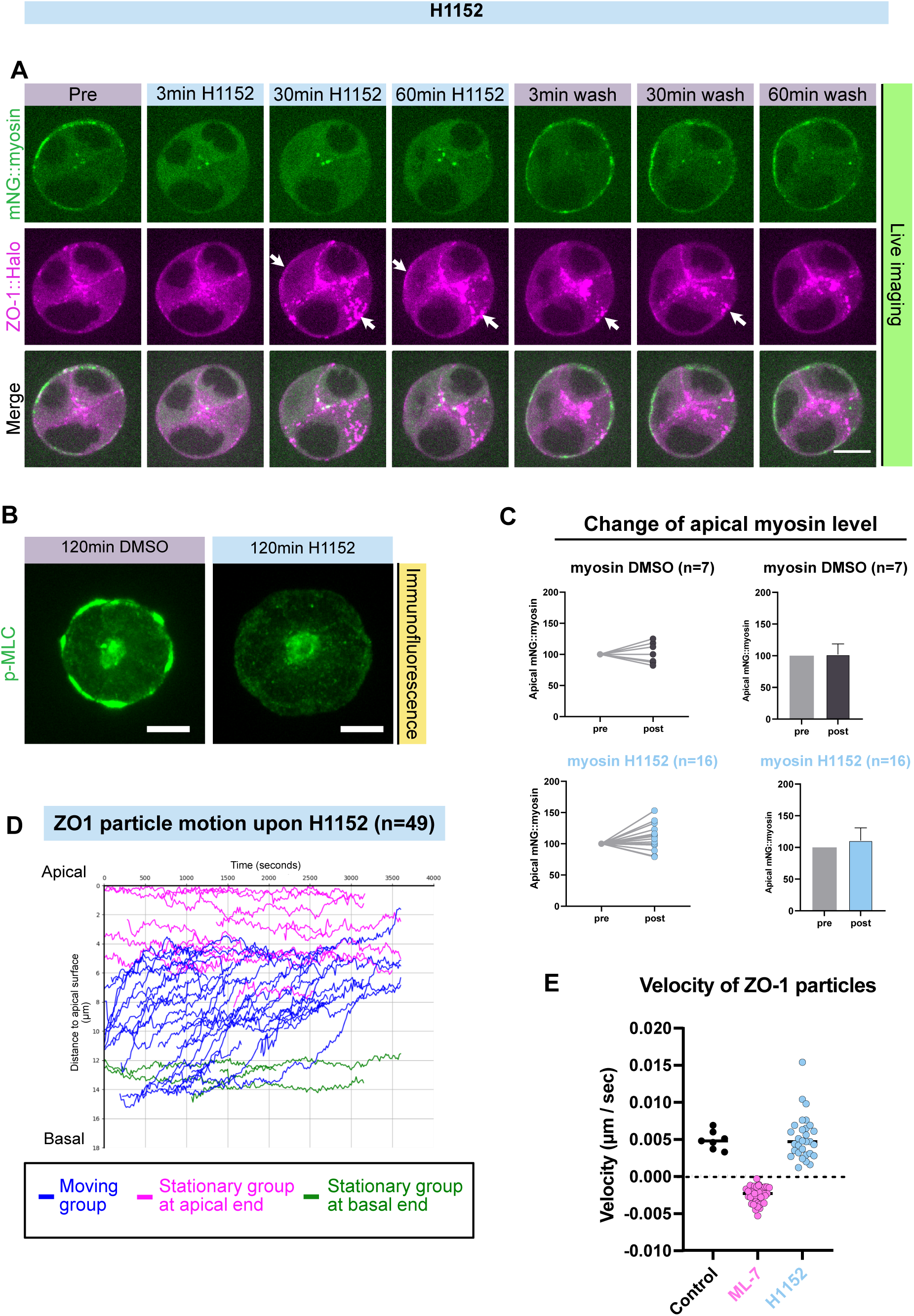
ROCK inhibition targets the basal myosin pool. (A) Confocal timelapse images showing live mNG::myosin and ZO-1::Halo in dual-tagged mESCs during the establishment of polarity. The images cover three stages of H1152 treatment: before treatment, after treatment, and following the wash-off. Arrows indicates the basal accumulation of ZO-1. Each image is a maximum-intensity projection of 3 planes spanning a total of 3 µm. Images were captured immediately (3 minutes) after the drug was administered or washed off. (B) Confocal images of fixed mESC spheroids stained with anti-pMLC 48 hours after plating. Each image is a maximum-intensity projection of 7 planes spanning a total of 9 µm. (C) Normalized apical mNG::myosin percentages at the midplane, measured pre-treatment, and 60 minutes after treatment. The top panel relates to DMSO (n = 7) treatment, the bottom to H1152 (n = 16), with all data adjusted to pre-treatment values; for non-normalized data, see Figure S5D. The left panel presents individual data points at the three stages; the right panel depicts average values with standard deviation. The small increase in apical myosin levels following H1152 treatment was marginally statistically significant (p = 0.044) when considering normalized data, but non-significant (p = 0.20) when considering the raw data (paired t-test; see Table S1 for exact p values). (D) Plot of position as a function of time for each ZO-1 particle. 49 ZO-1 particles from 3 different spheres were tracked during polarity establishment. The observation times were around 60 minutes following 60-minute H1152 treatment. There were 3 groups of trajectories: 14/49 stationary trajectories at the apical end (magenta), 4/49 stationary trajectories at the basal end (green), and 31/49 moving trajectories, which migrated from basal to apical (blue). (E) Scatter plot showing the velocity of ZO-1 particles under three conditions: control (n = 7), 60 minutes after ML-7 treatment (n = 35), and 60 minutes following H1152 treatment (n = 30). Control data, representing apical-moving particles, is sourced from Figure 1G-H; ML-7 data, showing basal-moving particles, is from Figure 2A-B. H1152 data with apical-moving particles is from Figure 5D. Scale bars represent 10 μm.

Importantly, ZO-1 particles continued to exhibit apical movement following ROCK inhibition (Figure 5D, Figure S5E, Movie 8). The velocity of this movement was unchanged compared to untreated cells (Figure 5E). However, the proportion of particles moving towards the apical surface increased to 63% in the group treated with H1152 (Figure 5D, Figure S5E), compared to 25% in the control group (Figure 1G & 1H, Figure S1B & S1C). In contrast, when MLCK was inhibited, 73% of ZO-1 particles moved in the opposite direction (Figure 2A-D, Figure S2C & S2D) at a slower speed (Figure 5E, Figure S5F). Taken together, these data suggest that ZO-1’s movement is a consequence of the dynamic balance between apical and basal myosin contractility, and that manipulating myosin activation through MLCK or ROCK inhibition can significantly alter ZO-1’s transportation within the cell. Under normal conditions, ZO-1 mainly moves apically, suggesting that a force imbalance generates apically directed cortical flow. Inhibiting MLCK disrupts the apical myosin flow, causing ZO-1 to move towards the basal side. On the other hand, ROCK inhibition primarily impacts the basal myosin pool, allowing apically directed flows to ensnare a larger proportion of ZO-1 particles.

Based on our earlier findings with ML-7 treatment, it was evident that apical myosin activity plays a role in organizing the actin cytoskeleton. This prompted us to investigate the effects of H1152 on the organization of F-actin. Despite changes in ZO-1 localization across various stages of H1152 treatment, F-actin consistently remained apical in all conditions (Figure S5B & S5C). The distinct outcomes observed with ML-7 and H1152 suggest that the apical, MLCK-dependent pool of myosin is primarily responsible for maintaining F-actin organization in epithelial cells.

### ROCK excludes polarity proteins from the basal domain

We were intrigued by the observation that although ZO-1 particles continued moving apically immediately after ROCK inhibition (Figures 5D-E and S5E-F), a pool of ZO-1 appeared to accumulate at the basal membrane following more prolonged H1152 treatment (Figures 5A, 6A and S5A-C). To determine if other polarity proteins behaved similarly, we examined the localization of PCX and aPKC along with ZO-1 in both live and fixed cells. Interestingly, all three of these proteins remained present at the apical domain following H1152 treatment, but they also formed ectopic basal patches that grew stronger over time (Figure 6B-F, Figure S6B-E, Movie 8). While ZO-1 and PCX formed patches at the basal surface, their apical pools remained largely intact. In the case of aPKC, basal patches were observed in live but not fixed cells (Figure 6C & 6E, Figure S6D & S6E). This might be due to the relatively high background and limited sensitivity of immunofluorescence staining with the anti-aPKCζ antibody, or alternatively, aPKCζ and aPKCι might be regulated differently. Regardless, these data indicate that ROCK activity is required on longer timescales to exclude apical proteins from the basal surface. Notably, the basal patches emerged slowly (30-60 minutes post-treatment, Figure 6F), while the disappearing of basal myosin was immediate (Figure 5A). This temporal difference suggests dual roles of ROCK: it acts acutely to sustain basal myosin activity, and over the long term, it prevents the establishment of apical protein patches at the basal surface. Together, these data imply that ROCK and MLCK act cooperatively, yet in distinct ways, in spatial regulation of actomyosin and polarity proteins during polarity establishment.

**Figure 6:**
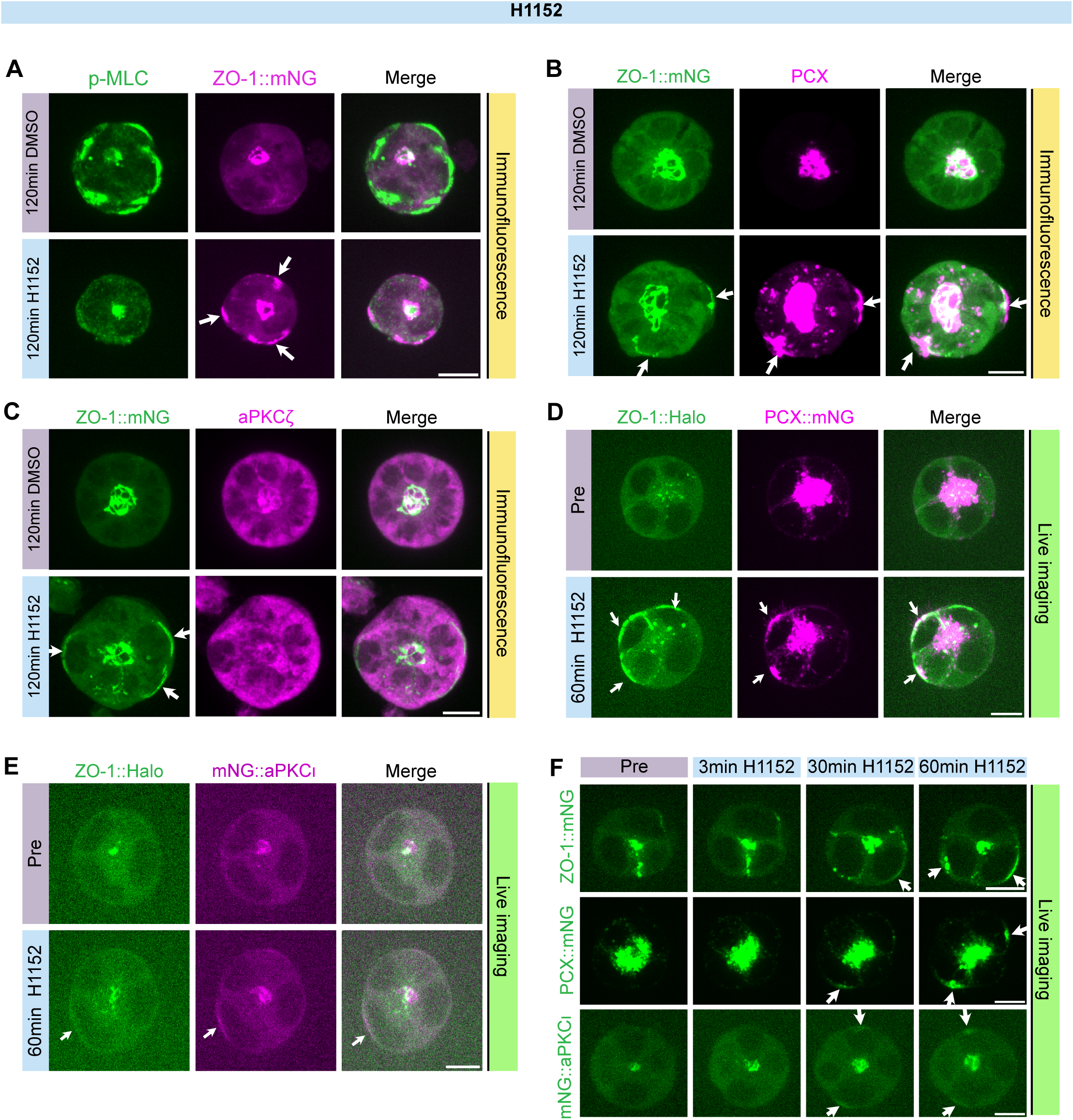
ROCK restricts the presence of polarity proteins from basal domain. (A) Confocal images of fixed ZO-1::mNG spheroids stained with the indicated antibody 48 hours after plating. The top panel shows spheroids treated with 120 minutes of DMSO, and the bottom panel with 120 minutes of H1152. Each image is a maximum-intensity projection of 7 planes spanning a total of 9 µm. (B) Confocal images of fixed ZO-1::mNG spheroids stained with the indicated antibody 48 hours after plating. The top panel shows spheroids treated with 120 minutes of DMSO, and the bottom panel with 120 minutes of H1152. Each image is a maximum-intensity projection of 7 planes spanning a total of 9 µm. (C) Confocal images of fixed ZO-1::mNG spheroids stained with the indicated antibody 48 hours after plating. The top panel shows spheroids treated with 120 minutes of DMSO, and the bottom panel with 120 minutes of H1152. Each image is a maximum-intensity projection of 7 planes spanning a total of 9 µm. (D) Confocal images showing live PCX::mNG and ZO-1::Halo in dual-tagged mESCs during the establishment of polarity. The images cover two stages of H1152 treatment: before treatment and post 60-minute H1152 treatment. Arrows indicates the basal accumulation of ZO-1. Each image is a maximum-intensity projection of 5 planes spanning a total of 6 µm. (E) Confocal images showing live mNG:: aPKCι and ZO-1::Halo in dual-tagged mESCs during the establishment of polarity. The images cover two stages of H1152 treatment: before treatment and post 60-minute H1152 treatment. Arrows indicates the basal accumulation of ZO-1. Each image is a maximum-intensity projection of 4 planes spanning a total of 4.5 µm. (F) Confocal timelapse images of ZO-1, PCX and aPKCι upon H1152 treatment during polarity establishment. Arrows indicates the basal accumulation of these polarity proteins. ZO-1 images are maximum-intensity projections of 4 planes covering 4.5 µm, while PCX and aPKCι images are projections of 5 planes spanning 6 µm. To note, images were captured immediately after the drug was administered. The initial image frame was taken approximately 3 minutes after the drug application. Scale bars represent 10 μm.

## Discussion

Cell polarity establishment is one of the earliest steps in embryonic development and is crucial for tissue formation. Central to this process is the targeted movement of polarity proteins to specific regions within cells. Using mESC spheroids as an epithelial polarity model, here we have shown that *de novo* establishment of the apical domain requires directional myosin-driven cortical flow. This flow transports the tight junction protein ZO-1, the adherens junction protein β-catenin, and likely many other components to establish the apical epithelial junction. Notably, we identified two separate pools of myosin, localized in the apical and basal domains. An apical, MLCK-dependent pool of myosin is required for apically directed flows and for polarity establishment and maintenance. A basal, ROCK-dependent pool of myosin appears to balance tension under normal conditions but leads to basal retraction of junctional components when MLCK is inhibited. Importantly, PCX and aPKC, which do not appear to flow along the lateral cell cortex, nevertheless require MLCK activity for their apical localization, indicating that polarized transport of junctional proteins is required for proper epithelial polarization.

The importance of cortical polarity flow for cell polarization is well established in the *C. elegans* zygote, a widely studied invertebrate model of cell polarity. In this system, anterior PARs proteins, including aPKC, are transported to the anterior side of the cell to establish the anterior domain (Munro et al., 2004; Cheeks et al., 2004; Motegi and Sugimoto, 2006; Motegi et al., 2011). In *C. elegans,* transport of aPKC by cortical flow relies on the oligomeric scaffold protein PAR-3, which recruits aPKC into cortical clusters whose stable membrane association allows them to be transported by cortical flow (Sailer et al., 2015; Chang and Dickinson, 2022; Dickinson et al., 2017; Illukkumbura et al., 2023). In mESC spheroids, we observed movement of a different scaffold protein, ZO-1, which is not present in *C. elegans* zygotes. More work will be required to determine whether ZO-1 in mammalian cells undergoes transport following similar principles as PAR-3 in *C. elegans,* but there are some interesting superficial similarities. Like PAR-3 (Lang et al., 2023), ZO-1 is a scaffold protein that can self-associate and that undergoes dynamic exchange between membrane and cytoplasmic pools (Shen et al., 2008; Beutel et al., 2019; Rouaud et al., 2020). Moreover, we observed colocalization between ZO-1 and aPKC, aligning with previous findings on MDCK cells(Suzuki et al., 2001) and suggesting that ZO-1 could play a role in recruiting aPKC to the plasma membrane. In H1152-treated cells, ZO-1 and aPKC were mislocalized together to the basal membrane, further supporting our conjecture of a physical association between these two proteins. However, in MDCK cells, ZO-1 knockout disrupted lumenogenesis without strongly affecting aPKC localization (Odenwald et al., 2016, 2018), so an alternative hypothesis is that PAR-3 itself, or another scaffold that is transported by flow, is responsible for polarizing both aPKC and ZO-1. Future experiments will aim to resolve this issue.

Our experiments showed that inhibition of MLCK by ML-7 interrupted ZO-1’s apical flow. After ML-7 removal, apical ZO-1 flow resumed, indicating its dependence on MLCK-driven myosin activity. Surprisingly, MLCK inhibition did not completely stop ZO-1 movement; instead, it reversed the direction. Retraction of ZO-1, observed upon MLCK inhibition, indicated that another factor is responsible for this reversed or basal actomyosin flow. Examining myosin distribution, we noted its presence in both apical and basal regions, consistent with earlier observations in mESCs (Shi et al., 2023) and human embryonic human pluripotent stem cell–derived cysts (Wang et al., 2021). We also observed distinct effects of inhibiting MLCK and ROCK on myosin activity and ZO-1 localization in cells. When MLCK was inhibited, there was a notable impact specifically on apical myosin leading to the relocation of ZO-1 towards the basal surface of the cell. In contrast, inhibiting ROCK selectively affected basal myosin, and allowed a larger amount of ZO-1 to flow towards the apical surface. These observations suggest that in typical conditions, forces are generated by both apical, MLCK-dependent and basal, ROCK-dependent myosin pools. We suspect that apical cortical tension is stronger, leading to flow of myosin that is directed predominantly towards the apical surface. Proper establishment of epithelial polarity, particularly the transportation of ZO-1, seems to hinge on this balance between apical and basal actomyosin tensions.

A basal function for ROCK is consistent with earlier work in other systems. This kinase has been noted to induce the formation of stress fibers and focal adhesion in fibroblasts, stimulated by ECM (Amano et al., 1997). Furthermore, during Drosophila oogenesis, both myosin and ROCK displayed a synchronized shift from the apical to basal domain, and ROCK positively influenced basal myosin intensity (Qin et al., 2018). More surprisingly, we found that ROCK inhibition led to an abnormal accumulation of ZO-1 basally, suggesting ROCK also prevents ZO-1’s basal localization. Basal stimulation of myosin and exclusion of apical proteins appear to be separate functions of ROCK, because loss of basal myosin happens immediately upon ROCK inhibition while ectopic basal localization of polarity proteins did not appear for 30-60 minutes. Although ROCK is best known for promoting myosin activity through MLC activation and myosin phosphatase inhibition, ROCK has diverse roles as a kinase and can target other substrates (Yoneda et al., 2005), including PAR-3 in *Drosopila* (Simões et al., 2010). ZO-1 has several sites matching the ROCK consensus motif (R/KXXS/T or R/KXS/T (Julian and Olson, 2014)), so it is possible that ZO-1 may be a direct ROCK substrate. Alternatively, ROCK activity may influence ZO-1 localization by impacting local cytoskeletal tension. Tension regulates a ZO-1 conformational transition from a folded to a stretched state which is crucial for its junctional location (Spadaro et al., 2017). Of course, ROCK could also regulate basal ZO-1 accumulation via an indirect mechanism. Further experiments are needed to elucidate the full spectrum of ROCK’s impact on apical polarity protein positioning.

In conclusion, our study elucidates the intricate dynamics of epithelial polarity establishment. We propose the opposite apical and basal actomyosin pools regulate epithelial polarization in primed mESCs.

## Materials and Methods

### Cell culture

Wild-type J1 (ATCC, SCRC-1010) and fluorescent knock-in mESCs were maintained in 2i/LIF (hereafter referred to as Naïve medium) at 37°C, 7% CO2 in gelatin-coated plates (Mulas et al., 2019). Naïve medium was made of N2B27 with 3 µM of GSK3β inhibitor (Sigma SML1046), 1 µM of MEK inhibitor (Sigma PZ0162), 100 U/mL of Leukemia Inhibitory Factor (Sigma ESG1106) and 1x Penicillin-Streptomycin (Thermo Fisher 15140122). N2B27 was a 1:1 mixture of DMEM/F12 (Sigma D6421) and Neurobasal (Fisher 21103049) supplemented with 0.5X B27 (Invitrogen 17504044), 1X N2 (homemade, see below), 50 μM β-mercaptoethanol (Sigma M3148-25ML), and 2 mM L-glutamine (Fisher 25030081). Homemade N2 contained DMEM/F12 (Sigma D6421), 2.5 mg/mL insulin (Sigma I9278), 10 mg/mL Apo-transferrin (Sigma T1147), 0.75% bovine albumin fraction V (Fisher 15260037), 1.98 μg/mL progesterone (Sigma P8783-1G), 1.6 mg/mL putrescine dihydrochloride (Sigma P5780-5G) and 0.518 μg/mL sodium selenite (Sigma S5261-10G). The final density for cell seeding was 15 X 10^4 cells/mL. Naïve medium was changed every day and mESCs were routinely passaged every 2-3 days. Mycoplasma testing (Southern Biotech 13100-01) was performed periodically.

For routine passaging of mESCs in gelatin-coated 10cm plates, Naïve medium was removed and 3.5 mL Accutase (Sigma SCR005) was added to cells and incubated at 37°C for 5 min. To dissociate mESCs into single cell suspension, cells were pipetted up and down 10-15 times. Then, the cell suspension was added to 10.5 mL of wash medium (500 mL of DMEM/F12 + 8 mL of bovine albumin fraction V) in a 15 mL tube. Cells were spun down for 5 min at 1000 rpm, and 1 mL of Naïve medium was used to resuspend cells. 10 µL cells were mixed with 10 μL of Trypan blue (Fisher 15250061) and counted using a cell counter (Invitrogen AMQAF1000). Finally, 150 X 10^6 cells with 10 mL of Naïve medium were seeded in a new gelatin-coated 10 cm plate.

### Fluorescent protein tagging

PCX::mNG mESCs were built by using the sleeping beauty transposon system(Mátés et al., 2009; Kowarz et al., 2015). pSB-ENN containing inverted terminal repeats (ITRs) and pCMV(CAT)T7-SB100 containing SB100X transposase were gifts from Jonghwan Kim (UT Austin). In the original pSB-ENN vector, EGFP flanked by ITRs was constitutively expressed. To switch this expression to PCX::mNG, we built the pSB-ENN-PCX-mNG vector by replacing the EGFP gene with the PCX::mNG gene in pSB-ENN vector. Then 950 ng pSB-ENN-PCX-mNG and 50 ng pCMV(CAT)T7-SB100 were mixed and diluted to 50 µL in Opti-MEM Medium (Gibco 31985062). 2 µL Lipofectamine 2000 (Fisher 11668027) was diluted to 50 µL in Opti-MEM Medium, then added to the 50 μL of DNA/Opti-MEM mixture and incubated at room temperature for 5 min. After incubation, the 100 μL DNA/Opti-MEM/Lipofectamine 2000 mixture was transferred to a gelatin-coated 12-well plate. 5 X 10^5 single cells in suspension per well were added to DNA/Opti-MEM/Lipofectamine 2000 mixture. Naïve medium was added up to 1 mL per well. The same procedures were performed with 50 μL of pure Opti-MEM Medium and 50 μL of diluted Lipofectamine 2000 as a negative control. We replaced the transfection mixture with 2-3 mL of fresh Naïve medium in the transfected cells on the next day. On day 3 after transfection, transfected cells were treated with Naïve medium+1 µg/mL Puromycin for 1 day, Naïve medium+0.5 µg/mL Puromycin for 1 day. PCX::mNG mESCs underwent verification through assessing cell viability and PCX distribution which was compared with the immunofluorescence result in wild-type cells to ensure that any findings were not due to potential issues from overexpression artifacts.

Other mESC lines were established by endogenous tagging utilizing CRISPR/Cas9-induced non-homologous end joining plus universal donors following methods previously published by our laboratory(Shi et al., 2023). Briefly, we employed a modular system comprising three plasmids: an FP donor vector, a frame selector Cas9/sgRNA vector, and a target Cas9/sgRNA vector. For tagging myosin, ZO-1, and aPKCι, we custom-designed target Cas9/sgRNA vectors for the Myh9, Tjp1, and Prkci genes. The other required vectors were already pre-made. These three plasmids were then introduced into mESCs using Lipofectamine 2000 (Thermo Fisher Scientific, 11668027). To isolate cells with successful tagging, drug selection and monoclonal picking were performed. The accuracy of the knock-in was confirmed through genotyping, Sanger sequencing, and analysis of protein distribution. Tables S2–S4 provide details on the cell lines, sgRNAs for CRISPR knock-in, and genotyping primers.

### Epithelial polarity establishment in 3D culture

To model polarity in 3D spheroid culture, mESCs were first primed in 2D. 30 X 10^4 cell mESCs were seeded with N2B27 (hereafter referred to as Naïve Pluripotency Exit (NPE) medium) in a 12-well gelatin-coated plate for 24 hours. On the second day, the primed mESCs were transferred into 3D setting to establish spheroids. Growth factor reduced Matrigel (Corning 356230) was used in 3D spheroid cultures. One well of a 15-well imaging plate (ibidi 81506) was covered with 1 µL of ice-cold Matrigel and incubated for 15 minutes at 37°C to allow the solidification of Matrigel. The primed mESCs were dissociated into single cell suspension with NPE medium, and cell density was adjusted to 10 X 10^4 cell/mL. Then 30 µL of cells (0.3 X 10^4 cell per well) were seeded on each Matrigel-coated well. When the cells had settled down to the Matrigel (15 minutes after seeding), the media were removed and replaced with 50 µL of NPE medium containing 5% Matrigel. Spheroids established epithelial polarity beginning approximately 24 hours after plating in 3D.

### Myosin inhibitor treatment

5 mg of ML-7 (Sigma I2764) was dissolved in 500 µL of DMSO, creating a stock solution with a concentration of 22 mM. 1 mg of H1152 (Fisher 24141) was dissolved in 51 µL of DMSO to prepare a stock solution with a concentration of 50 mM. For storage, both ML-7 and H1152 solution were aliquoted into 2 µL per vial and stored at -80°C. To achieve a working concentration of 40 µM ML-7 for drug perturbation experiments (Connell and Helfman, 2006), 2 µL of the 22 mM stock solution was diluted in 1100 µL of NPE medium containing 5% Matrigel. To prepare a working concentration of 50 µM H1152 for drug perturbation (Zenker et al., 2018), 1 µL of the 50 mM stock solution was diluted in 999 µL of NPE medium containing 5% Matrigel. DMSO was also diluted in the same ratio to serve as a control.

For drug perturbation, the existing medium was removed from the cells, and drug-containing medium was added for 1-2 hour. To conduct wash-off experiments, the drug-containing medium was removed, and the cells were washed twice with wash medium (comprising 500 mL of DMEM/F12 + 8 mL of bovine albumin fraction V). Then, NPE medium containing 5% Matrigel was added.

Table S5 lists drugs used in this study and their sources.

### Immunofluorescence

Medium was removed and cells were fixed with 4% PBS-paraformaldehyde (Thermo Fisher J60401-AK) for 20 minutes at room temperature. Then PBS was used to wash three times (5 minutes each). Then we incubated the fixed samples with permeabilization buffer, composed of 0.3% Triton X-100 and 0.1M glycine in PBS for 20 minutes at room temperature. PBST buffer with 0.1% Tween in PBS was used to wash three times (10 minutes each with rocking). To block spheroids, we incubated samples in blocking buffer (10% serum in PBST) for 2 hours at room temperature. Primary antibodies were incubated overnight at 4°C on a rocker, followed by washing in PBST (3 times for 1 hour each). Secondary antibodies were applied for 1 hour in the dark at room temperature, followed by washing in PBST (3 times for 1 hour each) and then PBS (twice for 10 minutes each). We aspirated the PBS and mounted slides with ProLong Mountant with DAPI (Thermo Fisher P36941-2ml) for imaging. Table S6 provides a list of antibodies used in this study, along with the respective dilution ratios for each.

### Dye labeling for live imaging

To image cells with HaloTag, HaloTag ligand dye JF_646_ (gift from Luke Lavis) was used. It was firstly dissolved in acetonitrile to 1 mM and aliquoted into 2 µL in PCR tubes. The dye was dried with a vacuum centrifuge for storage at -20°C in a desiccator. Before imaging, we dissolved one aliquot in 2 µL of DMSO to reconstitute 1 mM dye. Then the dye was added to NPE medium to a final concentration of 0.037 µM. Before imaging, we incubated cells with 0.037 µM JF_646_ for at least 30 minutes in a 15-well imaging plate at 37°C. After the incubation, we removed JF_646_ and washed the spheroids with wash medium (500 mL of DMEM/F12 + 8 mL of bovine albumin fraction V) 3 times. Then 50 µL of NPE medium containing 5% Matrigel were added.

Cell tracing dyes including SPY650-FastAct (CY-SC505) and SPY650-DNA (Cytoskeleton CY-SC501) were dissolved in 50 µL of fresh DMSO to reconstitute 1000X. Then they were aliquoted into 5 µl in PCR tubes, protected from light and stored at -80°C. Before imaging, we incubated cells with 1X cell tracing dye with NPE medium containing 5% Matrigel overnight in a 15-well imaging plate at 37°C until cells were ready for imaging. There was no wash step after dye incubation because all dyes were fluorogenic and produced low background fluorescence.

Table S7 lists dyes used for cell labeling in this study.

### Confocal imaging

For fixed samples, images were acquired using one of two microscopes. The images shown in Figure 1B-C were acquired using a Nikon Eclipse Ti-2 microscope equipped with a 60X, 1.4 NA objective; a Photometrics PrimeBSI camera; an OptoSpin filter wheel (CAIRN Research, Kent, England), and an vt-iSIM super-resolution confocal scan head (VisiTech international, Sunderland, UK). All other images were acquired using a Nikon Eclipse Ti-2 microscope equipped with a 60X, 1.4 NA objective; an 89 North LDI-7 laser diode illuminator; a Photometrics Prime95B camera; and a Crest X-Light V3 spinning disk confocal head. mESCs were imaged by multiple z-planes with a step of 1.5 µm.

Live sample images were acquired using same spinning disk confocal setup described above for fixed samples. mESCs were seeded in 15-well imaging plate and incubated in a tissue culture incubator until imaging was initiated. For observing the establishment of epithelial polarity in 3D culture, cells were incubated for a minimum of 24 hours. Prior to imaging, a humidifier was filled with water, and a gas mixer (Okolab 2GF-MIXER) with an air pump, 100% CO2, and a temperature controller (Okolab H401-T-CONTROLLER) were activated to pre-equilibrate the imaging chamber (Okolab H201-NIKON-TI-S-ER) to the correct conditions: 7% CO2, 37°C, and optimal humidity for live cells. Snapshot imaging of mESCs was performed across multiple z-planes with a step of 1.5 µm at specific times. For particle tracking, fast time-lapse imaging was acquired every 15 or 30 seconds in 2-4 z-planes close to the mid-plane, with a step of 1.5 µm, spanning 20-70 minutes. mNG, SPY650-FastAct, SPY650-DNA, and HaloTag-JF646 were excited using the appropriate lines of the LDI-7 illuminator.

For figure preparation, images were cropped and rotated using FIJI software, and adjustments were made to brightness and contrast to enhance signal visibility. No additional image manipulations were performed. Table S8 lists all software used for data analysis in this study.

### Particle tracking and data analysis

To track ZO-1 particles, the maximum z projection of the time-lapse movie, focusing on the ZO-1 channel, was processed using the TrackMate FIJI plugin (Ershov et al., 2022). The detector utilized was the Difference of Gaussian (DoG), and an estimated object diameter was set at 0.9 µm. A quality threshold was established at 1.301, and pre-processing and sub-pixel localization were enabled. Spots were color-coded based on track ID for ease of identification. The Linear Assignment Problem (LAP) tracker (Jaqaman et al., 2008) was selected as the tracker, and frame-to-frame linking was set with a maximum distance of 1 µm. Track segment gap closing was configured to 2 µm, allowing for a maximum gap of 2 frames. Filters were implemented on tracks, focusing on the number of spots in a track and track mean quality, to systematically exclude artificial and low-quality tracks from the analysis.

Upon obtaining spots and tracks information from multiple spheres, which represent the movement of ZO-1 particles in the movies, we manually defined the apical (center) and basal (peripheral) surfaces of the spheres. Subsequently, all sphere radii were normalized to a value of 1 to allow combining data from multiple spheroids. The normalized data were plotted on a trajectory graph, with the apical end positioned at the center and the basal end situated on the circle with a radius equal to 1, as illustrated in Figures 1G, S1B, 2C and S2C. Trajectories were categorized and color-coded based on their respective motion patterns to facilitate clear and distinct visualization of the various movement pathways of the ZO-1 particles.

Additionally, we plotted the position as a function of time for each ZO-1 particle, presented in Figures 1H, S1C, 2D, S2D, 5D and S5E. In these graphs, the position was measured as the distance from the apical surface, with the apical surface itself defined as the zero point.

For the velocity analysis, we selected raw data representing apical-moving particles from control data in Figure 1G-H (n = 8), basal-moving particles from ML-7 data in Figure 2A-B (n = 38), and apical-moving particles from H1152 data in Figure 5D (n = 31). This data was then refined by first eliminating plateaus in the trajectories, followed by excluding tracks with short durations (less than 300 seconds: 1 from the control group, 3 from the ML-7 group, and 1 from the H1152 group). We then created scatter plots for each tracked ZO-1 particle, plotting position as a function of time (Figure S5F). To best represent the data, we applied a linear regression equation: y=mx+b, where y represents position, x time, m the velocity (slope of the line), and b the y-intercept.

### Measurement of apical protein level

The process for quantifying the percentage of various apical proteins at the midplane is detailed in Figure S2E. We used Fiji software to measure the intensity levels. The calculation for the intensity at the apical domain (Int_apical) is as follows:

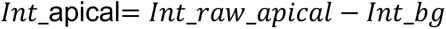

Similarly, the intensity of the entire sphere (Int_total) is calculated as below:

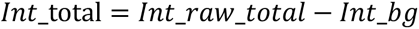

Subsequently, the percentage of apical intensity (%apical) is calculated with the following equation:

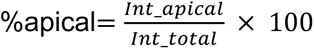

Where: Int denotes the integrated density. Int_raw_apical or Int_raw_basal is the integrated density measured within a circle proximate to the apical or basal domain, respectively. Int_bg is the integrated density of the background measured from the same region of interest (ROI) as apical or basal circles.

Raw percentage data is displayed in Figures S2F, S3C-D, and S5D. For direct comparison, post-treatment and post-wash-off data were normalized to pre-treatment levels in Figures 2F, 3F-G, and 5C.

All statistical analyses were performed using Graphpad Prism. Tests used are listed in each figure legend, and exact p values are listed in Table S1.

## Data Availability Statement

All primary data supporting this manuscript will be openly available in the Texas Data Repository upon manuscript acceptance for publication. The data will be publicly accessible at https://doi.org/10.18738/T8/YTOBXT.

## Acknowledgements

We thank members of the Dickinson lab for helpful discussions and comments on the manuscript, and Lea Jinks for technical assistance. This work was supported by NSF MCB 2237451 (DJD).

---

**Figure S1.**
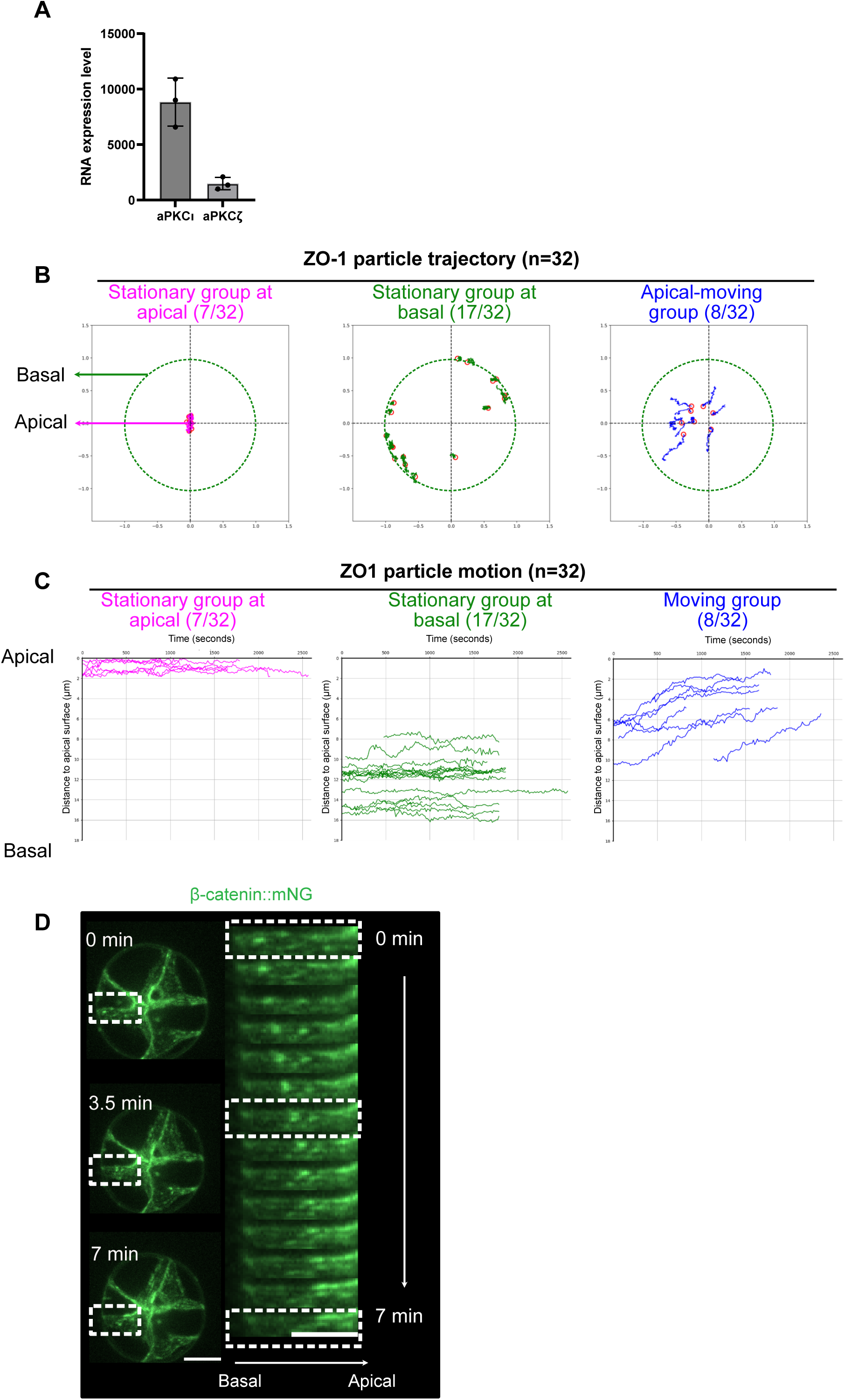
(A) RNA expression of two aPKC isoforms, aPKCι and aPKCζ using data from the published RNA sequencing of 3D spheroids in primed mESCs (GSE141773). We tagged aPKCι rather than aPKCζ due to its higher expression in primed mESCs. (B) Trajectories of 32 ZO-1 particles from 4 different spheres during polarity establishment. The observation times ranged from 27.5 to 42.75 minutes. There were 3 groups of trajectories: 7/32 stationary trajectories at the apical end (magenta), 17/32 stationary trajectories at the basal end (green), and 8/32 moving trajectories, which migrated from basal to apical (blue). Red circles indicate the ends of each trajectory. The basal surface is represented by a green dash circle (normalized radius = 1) and the apical surface is at the center. (C) Plot of position as a function of time for each tracked ZO-1 particle represented in (B). (D) Confocal live images showing the movement of β-catenin during polarity establishment. The box on the left is magnified on the right. Each image is a maximum-intensity projection of 5 planes spanning a total of 6 µm.

**Figure S2.**
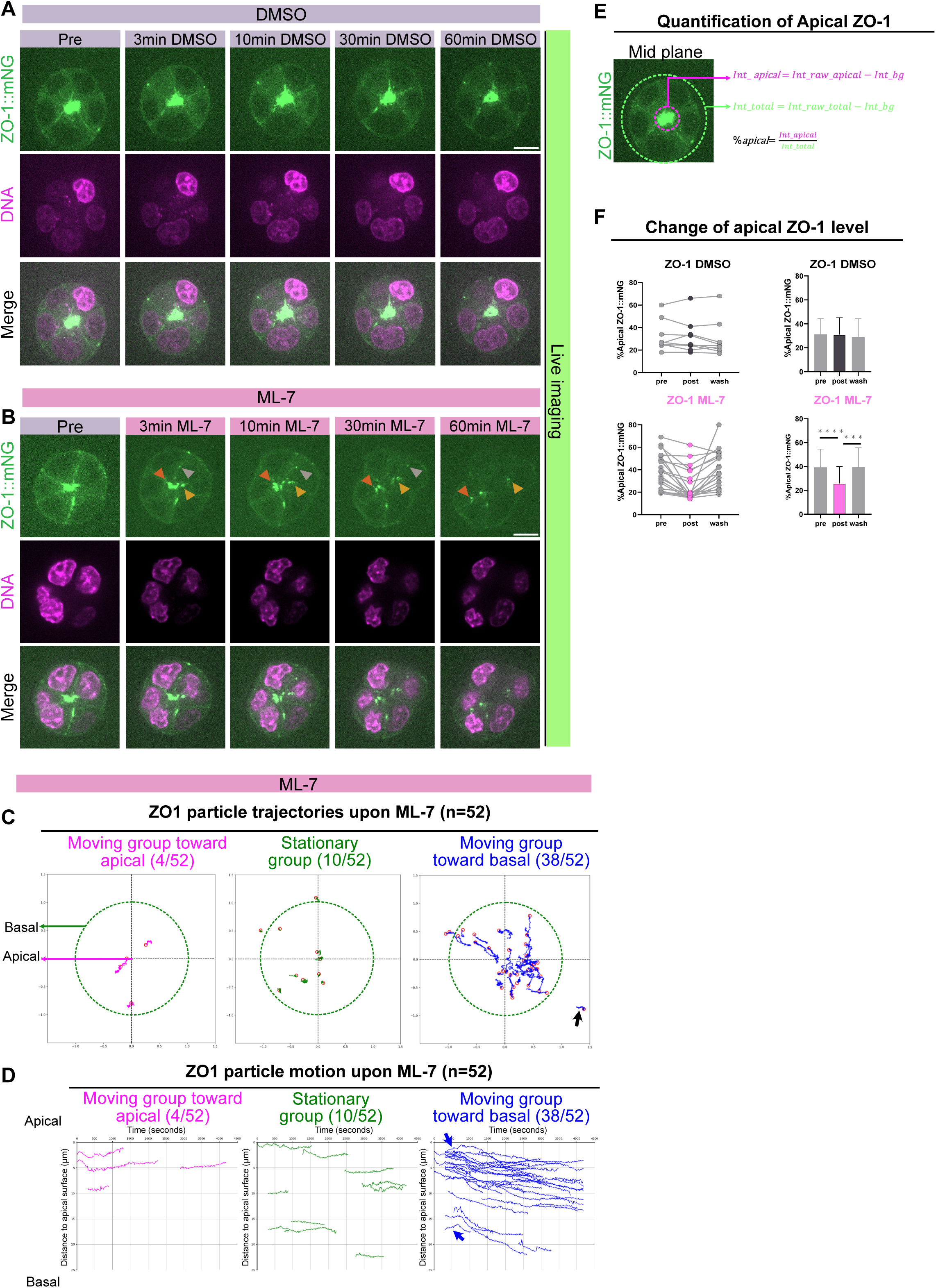
(A) Confocal images of ZO-1 and nuclei pre- and post-DMSO treatment. Each image is a maximum-intensity projection of 3 planes spanning a total of 3 µm. (B) Confocal images of ZO-1 and nuclei pre- and post-ML-7 treatment. Each image is a maximum-intensity projection of 3 planes spanning a total of 3 µm. (C) Trajectories of 52 ZO-1 particles from 3 different spheres during polarity establishment. The observation times were 70 minutes following 60-minute ML-7 treatment. There are 3 groups of trajectories: 38/52 was basal-moving group which migrated from apical to basal towards (blue), 4/52 was apical-moving group (magenta) and 10/52 was stationary group (green). The red circles indicate the ends of each trajectory. The basal surface is represented by a green dash circle (normalized radius = 1) and the apical surface is at the center. The black arrow represents one basal-moving trajectory outside the original basal surface due to a distorted shape (radius > 1). (D) Plot of position as a function of time for each tracked ZO-1 particle represented in (C). Two blue arrows depict trajectories moving from basal to apical, then retracting back to basal. (E) Illustration of quantification to assess the percentage of apical ZO-1::mNG at the midplane. The intensity in the apical domain (Int_apical) is determined by subtracting the background intensity from the raw intensity measured within the small magenta circle. Similarly, the intensity of the entire sphere (Int_total) is calculated by subtracting the background intensity from the raw intensity measured within the larger green circle. Both background intensities are measured from the same region of interest (ROI) as their respective circles. The percentage of apical intensity (%apical) is then calculated by dividing Int_apical by Int_total. (F) Raw apical ZO-1::mNG percentages at the midplane, measured pre-treatment, 60 minutes after treatment, and 60 minutes post-wash. The top panel relates to DMSO (n = 10) treatment, the bottom to ML-7 (n = 20). The left panel presents individual data points at the three stages; the right panel depicts average values with standard deviation. Asterisks indicate significant changes (paired t-test; see Table S1 for exact p values). Scale bar: 10 μm.

**Figure S3.**
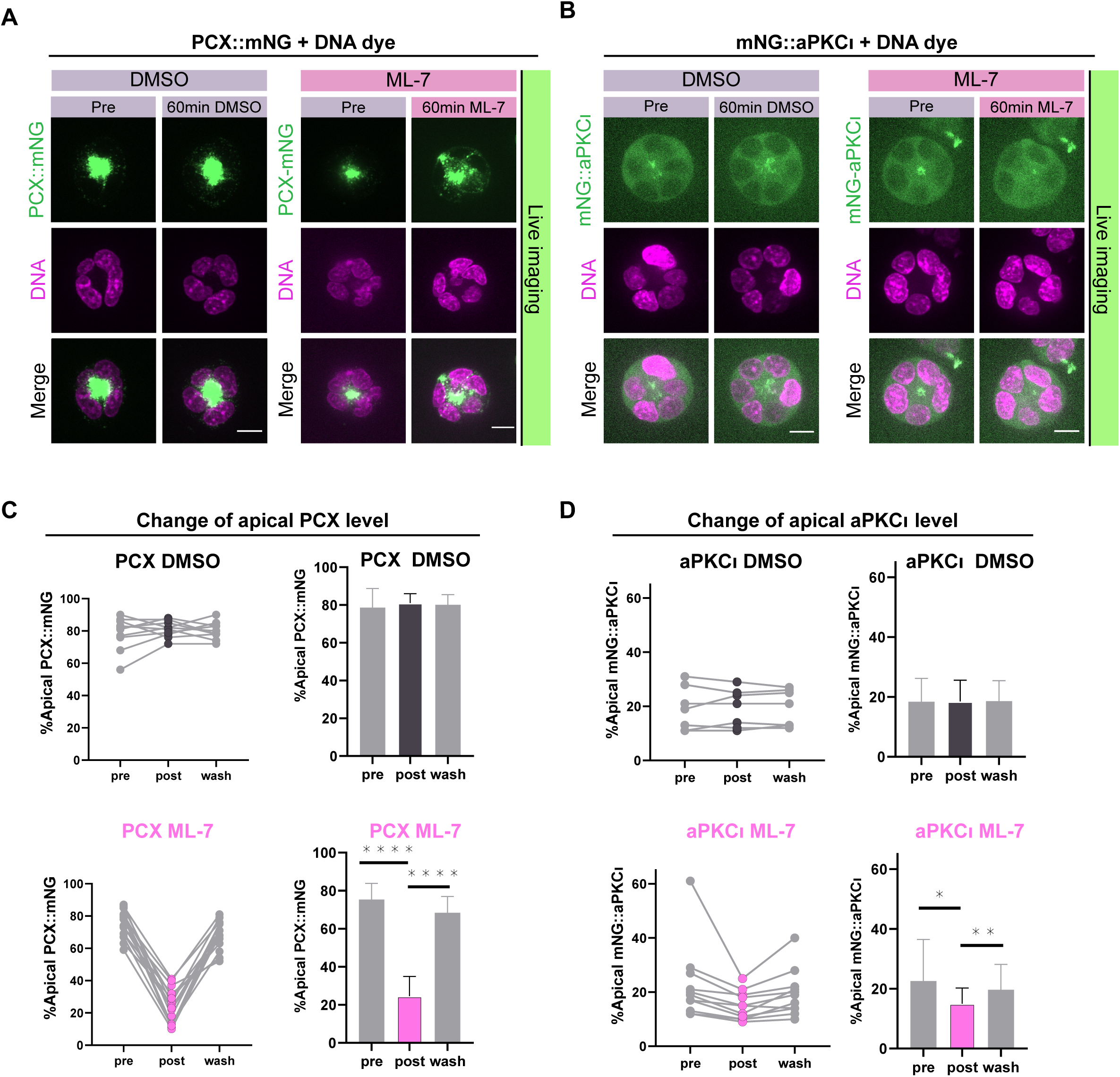
(A) Confocal images of PCX and nuclei before and after 60-minute DMSO or ML-7 treatment. Each image is a maximum-intensity projection of 5 planes spanning a total of 6 µm. (B) Confocal images of aPKCι and nuclei before and after DMSO or ML-7 treatment. Each image is a maximum-intensity projection of 5 planes spanning a total of 6 µm. (C) Raw apical PCX::mNG percentages at the midplane, measured pre-treatment, 60 minutes after treatment, and 60 minutes post-wash. The top panel relates to DMSO (n = 10) treatment, the bottom to ML-7 (n = 15). The left panel presents individual data points at the three stages; the right panel depicts average values with standard deviation. Significant changes are marked by asterisks (paired t-test; see Table S1 for exact p values). (D) Raw apical mNG::aPKCι percentages at the midplane, measured pre-treatment, 60 minutes after treatment, and 60 minutes post-wash. The top panel relates to DMSO (n = 8) treatment, the bottom to ML-7 (n = 11). The left panel presents individual data points at the three stages; the right panel depicts average values with standard deviation. Significant changes are marked by asterisks (paired t-test; see Table S1 for exact p values). Scale bar: 10 μm.

**Figure S4.**
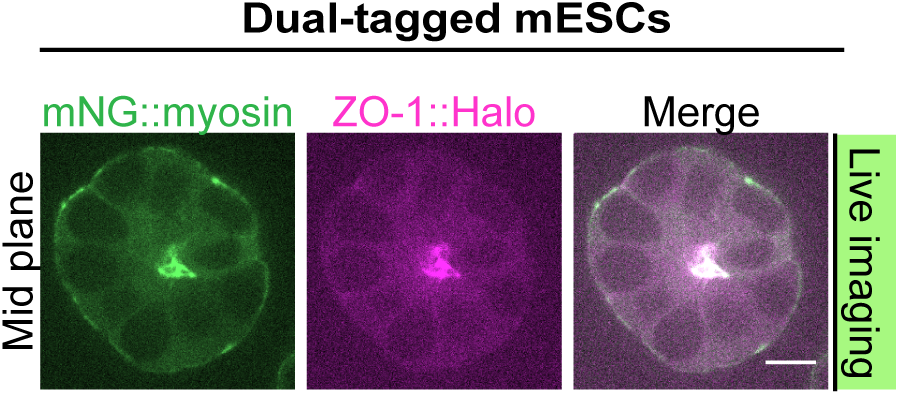
Mid-plane images of mNG::myosin and ZO-1::Halo at the midplane in dual-tagged mESCs 53 hours after plating. Scale bar: 10 μm.

**Figure S5.**
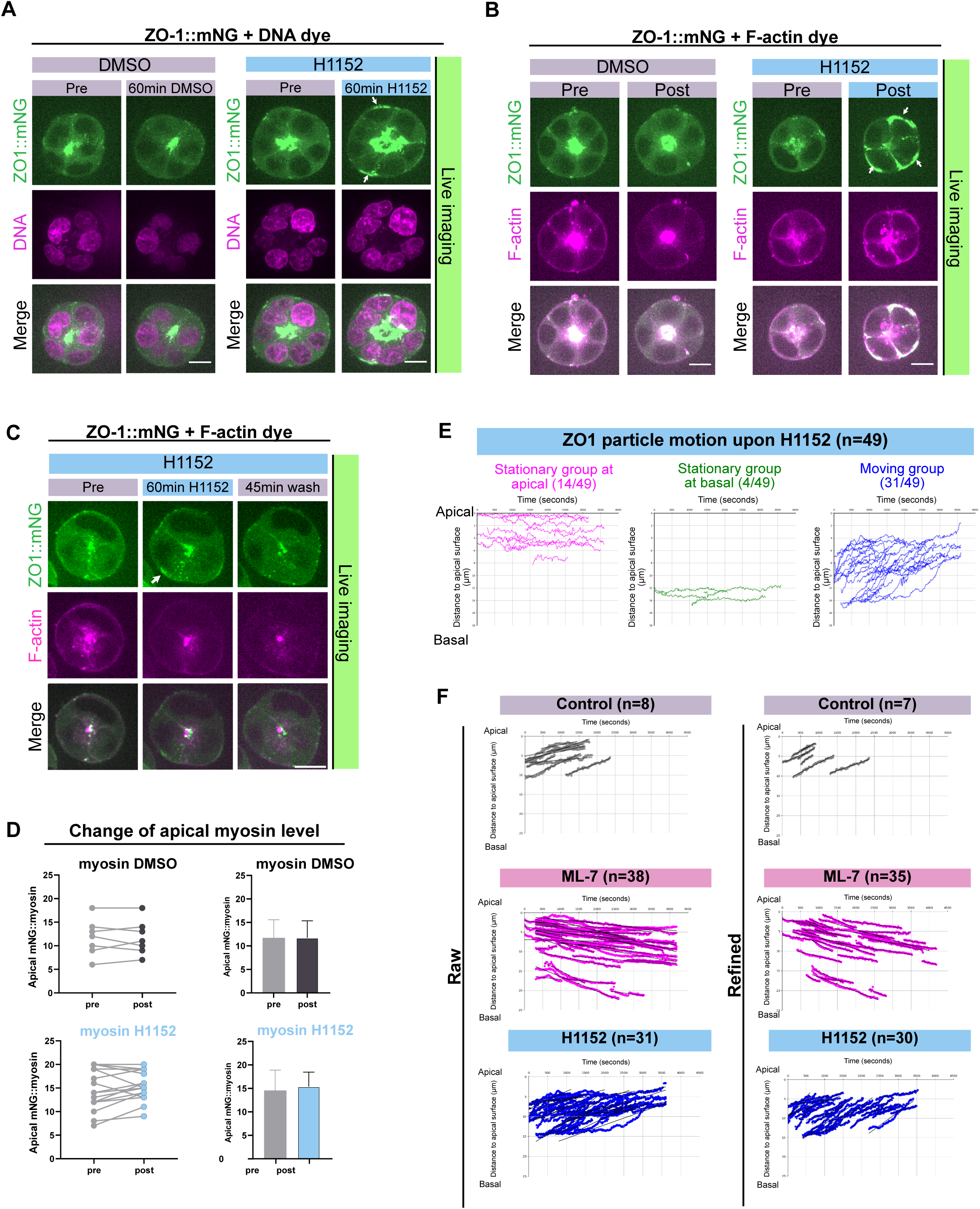
(A) Confocal images of ZO-1 and nuclei before and after 60-minute DMSO or H1152 treatment. Each image is a maximum-intensity projection of 5 planes spanning a total of 6 µm. (B) Confocal images of ZO-1 and F-actin before and after DMSO or H1152 treatment. Each image is a maximum-intensity projection of 5 planes spanning a total of 6 µm. (C) Confocal timelapse images showing ZO-1::mNG and F-actin during the establishment of polarity. The images cover three stages of H1152 treatment: before treatment, after 60-minute treatment, and following 45-minute wash-off. Arrows indicates the basal accumulation of ZO-1. Each image is a maximum-intensity projection of 3 planes spanning a total of 3 µm. (D) Raw apical mNG::myosin percentages at the midplane, measured pre-treatment, and 60 minutes after treatment. The top panel relates to DMSO (n = 7) treatment, the bottom to H1152 (n = 16). The left panel presents individual data points at the three stages; the right panel depicts average values with standard deviation. Significant changes are marked by asterisks (paired t-test; see Table S1 for exact p values). (E) Plot of position as a function of time for each ZO-1 particle. 49 ZO-1 particles from 3 different spheres were tracked during polarity establishment. The observation times were around 60 minutes following 60-minute H1152 treatment. There were 3 groups of trajectories: 14/49 stationary trajectories at the apical end (magenta), 4/49 stationary trajectories at the basal end (green), and 31/49 moving trajectories, which migrated from basal to apical (blue). (F) Plot of position as a function of time for each tracked ZO-1 particle with linear trends marked by black lines at 15-second intervals. Control data, representing apical-moving particles, is sourced from Figure 1G-H; ML-7 data, showing basal-moving particles, is from Figure 2A-B. H1152 data with apical-moving particles is from Figure 5D. The left panel displays raw data. The right panel shows refined data, achieved by first removing plateaus in the trajectories and then excluding those with short-duration tracks (tracking time < 300 seconds: 1 from the control group, 3 from the ML-7 group, 1 from the H1152 group). Scale bar: 10 μm.

**Figure S6.**
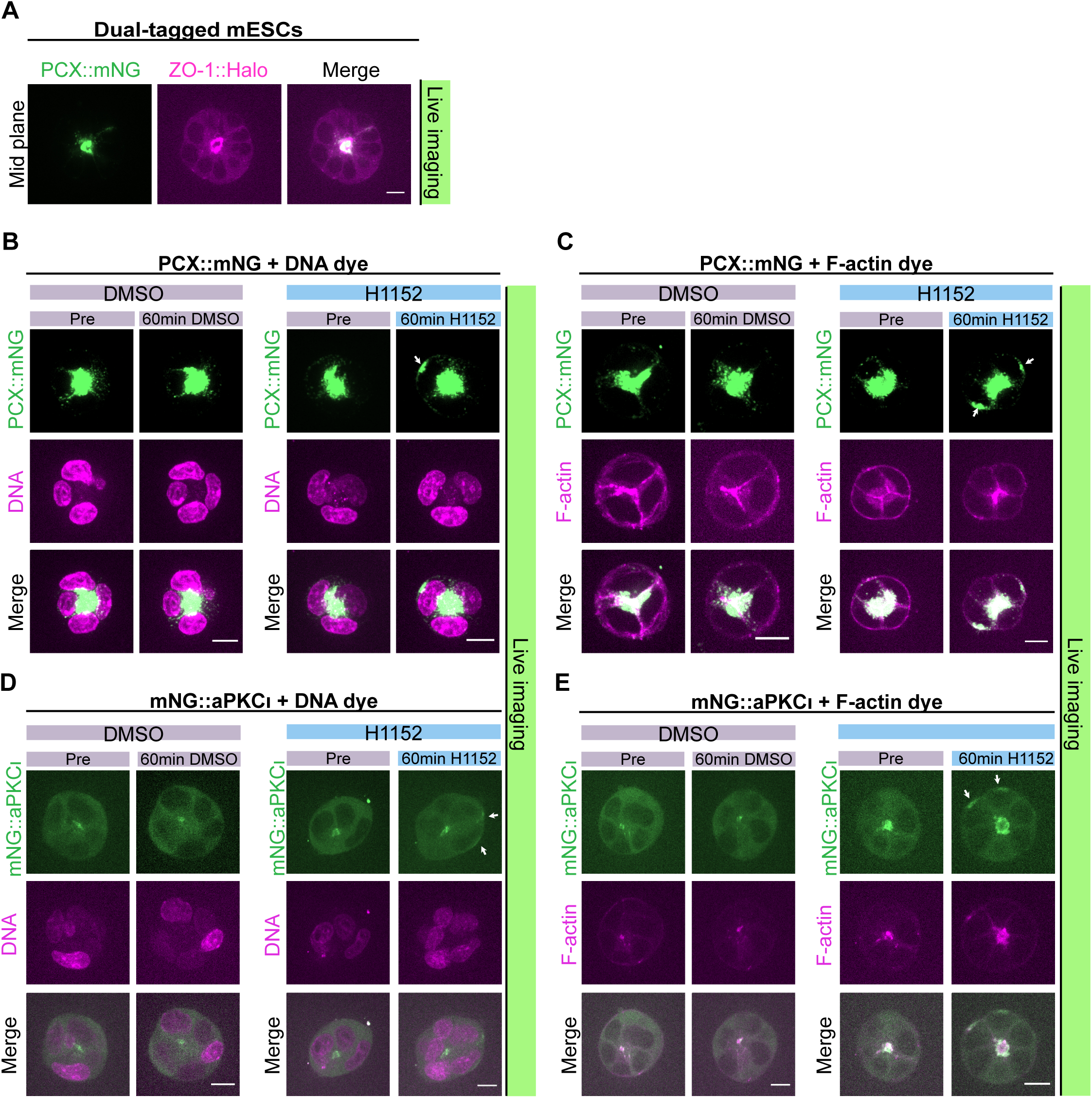
(A) Confocal images of PCX::mNG and ZO-1::Halo at the midplane in dual-tagged mESCs 60 hours after plating. (B) Confocal images showing live PCX and nuclei in dual-tagged mESCs during the establishment of polarity. The images cover two stages of H1152 treatment: before treatment and post 60-minute H1152 treatment. Arrows indicates the basal accumulation of ZO-1. Each image is a maximum-intensity projection of 4 planes spanning a total of 4.5 µm. (C) Confocal images showing live PCX and F-actin during the establishment of polarity. The images cover two stages of H1152 treatment: before treatment and post 60-minute H1152 treatment. Arrows indicates the basal accumulation of ZO-1. Each image is a maximum-intensity projection of 5 planes spanning a total of 6 µm. (D) Confocal images showing live aPKCι and nuclei during the establishment of polarity. The images cover two stages of H1152 treatment: before treatment and post 60-minute H1152 treatment. Arrows indicates the basal accumulation of ZO-1. Each image is a maximum-intensity projection of 5 planes spanning a total of 6 µm. (E) Confocal images showing live aPKCι and F-actin during the establishment of polarity. The images cover two stages of H1152 treatment: before treatment and post 60-minute H1152 treatment. Arrows indicates the basal accumulation of ZO-1. Each image is a maximum-intensity projection of 5 planes spanning a total of 6 µm. Scale bar: 10 μm.

**Table S1.** Results of Statistical Tests.

| Figure | Comparison | p value |
| --- | --- | --- |
| 2F | Pre vs. post DMSO | 0.77 |
| 2F | Post DMSO vs. washout | 0.063 |
| 2F | Pre vs. post ML-7 | <0.0001 |
| 2F | Post ML-7 vs. washout | 0.0002 |
| S2F | Pre vs. post DMSO | 0.90 |
| S2F | Post DMSO vs. washout | 0.11 |
| S2F | Pre vs. post ML-7 | <0.0001 |
| S2F | Post ML-7 vs. washout | 0.0004 |
| 3F | Pre vs. post DMSO | 0.30 |
| 3F | Post DMSO vs. washout | 0.90 |
| 3F | Pre vs. post ML-7 | <0.0001 |
| 3F | Post ML-7 vs. washout | <0.0001 |
| 3G | Pre vs. post DMSO | 0.96 |
| 3G | Post DMSO vs. washout | 0.40 |
| 3G | Pre vs. post ML-7 | <0.0001 |
| 3G | Post ML-7 vs. washout | 0.0015 |
| S3C | Pre vs. post DMSO | 0.36 |
| S3C | Post DMSO vs. washout | 0.65 |
| S3C | Pre vs. post ML-7 | <0.0001 |
| S3C | Post ML-7 vs. washout | <0.0001 |
| S3D | Pre vs. post DMSO | 0.90 |
| S3D | Post DMSO vs. washout | 0.78 |
| S3D | Pre vs. post ML-7 | 0.028 |
| S3D | Post ML-7 vs. washout | 0.0062 |
| 5C | Pre vs. post DMSO | 0.78 |
| 5C | Pre vs. post H1152 | 0.044 |
| S5D | Pre vs. post DMSO | >0.9999 |
| S5D | Pre vs. post H1152 | 0.20 |

**Table S2.** Cell lines.

| Cell line | Source | Identifier |
| --- | --- | --- |
| J1 Wild-type | ATCC | SCRC-1010 |
| ZO-1::mNG | (Shi et al., 2023) | N/A |
| mNG::myosin | (Shi et al., 2023) | N/A |
| β-catenin::mNG | (Shi et al., 2023) | N/A |
| PCX::mNG | This work | N/A |
| mNG::aPKCι | This work | N/A |
| mNG::myosin; ZO-1::Halo | This work | N/A |
| PCX::mNG; ZO-1::Halo | This work | N/A |
| mNG::aPKCι; ZO-1::Halo | This work | N/A |

**Table S3.**
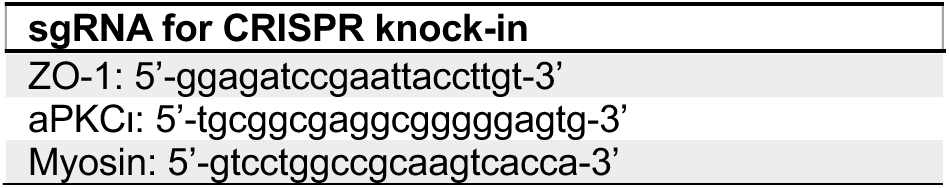
sgRNAs for CRISPR knock-in.

**Table S4.**
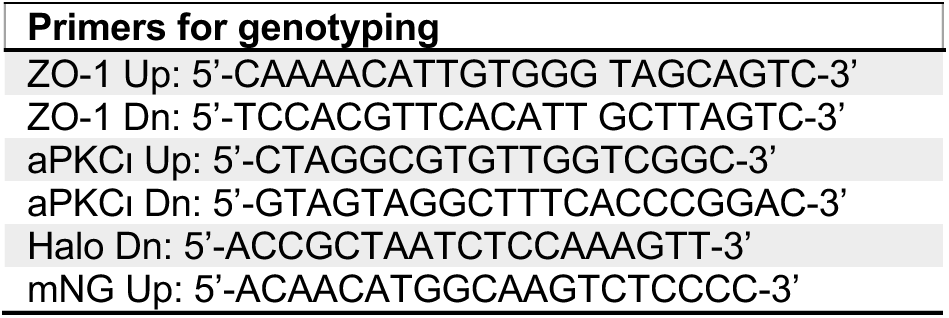
Primers for genotyping.

**Table S5.** Myosin inhibitors.

| Myosin inhibitors | Source | Identifier |
| --- | --- | --- |
| ML-7 | Sigma | I2764 |
| H 1152 DIHYDROCHLORIDE | Fisher | 24141 |

**Table S6.** Antibodies.

| Antibody | Source | Identifier |
| --- | --- | --- |
| Rabbit p-MLC (Ser19), 1:100 dilution | Fisher | PIPA517726 |
| Rabbit PAR-3, 1:200 dilution | Millipore | 07-330 |
| Rabbit Rab11a, 1:100 dilution | Cell Signaling Technology | 2413 |
| Rat E-cadherin, 1:200 dilution | Fisher | 13-1900 |
| Rat PCX, 1:250 dilution | Fisher | MAB1556 |
| Mouse ZO-1, 1:100 dilution | Fisher | 33-9100 |
| Mouse aPKCζ, 1:100 dilution | Santa Cruz Biotechnology | sc-17781 |
| Alexa Fluor 568 Donkey anti-Mouse, 1:500 dilution | Fisher | A10037 |
| Alexa Fluor 647 Donkey anti-Mouse, 1:500 dilution | Fisher | A31571 |
| Alexa Fluor 568 Donkey anti-Rabbit, 1:500 dilution | Fisher | A10042 |
| Alexa Fluor 647 Donkey anti-Rabbit, 1:500 dilution | Fisher | A-31573 |
| Alexa Fluor 488 Donkey anti-Rat, 1:500 dilution | Fisher | A-21208 |
| Alexa Fluor 594 Donkey anti-Rat, 1:500 dilution | Fisher | A21209 |
| Alexa Fluor 488 Phalloidin, 1:400 dilution | Fisher | A12379 |
| Alexa Fluor 555 Phalloidin, 1:400 dilution | Fisher | A34055 |
| Alexa Fluor 647 Phalloidin, 1:400 dilution | Fisher | A22287 |

**Table S7.**
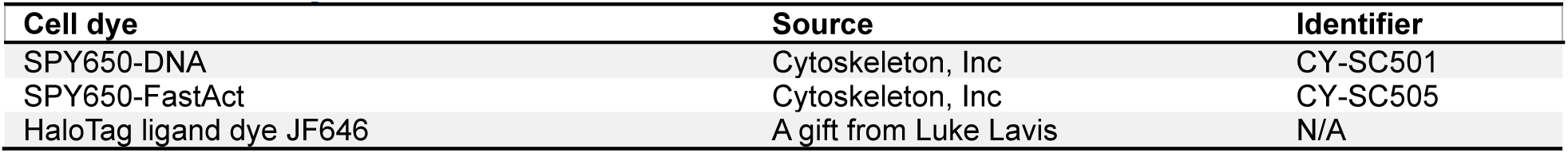
Cell dye.

| Cell dye | Source | Identifier |
| --- | --- | --- |
| SPY650-DNA | Cytoskeleton, Inc | CY-SC501 |
| SPY650-FastAct | Cytoskeleton, Inc | CY-SC505 |
| HaloTag ligand dye JF646 | A gift from Luke Lavis | N/A |

**Table S8.**
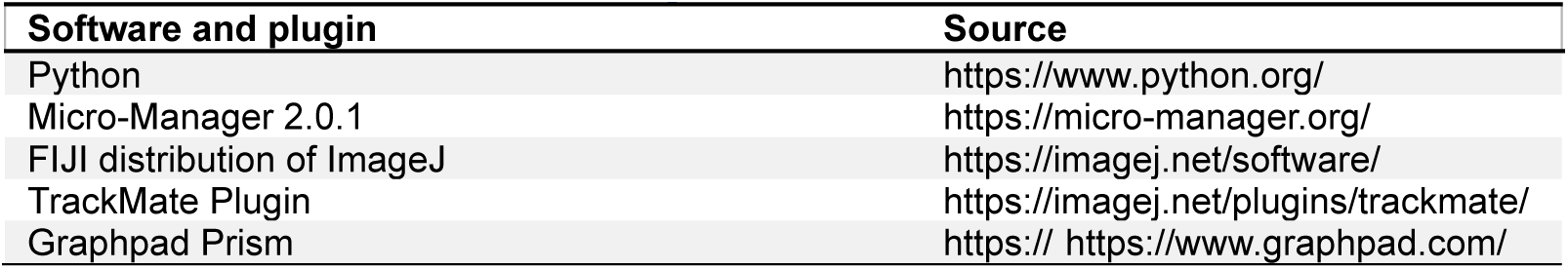
Software and plugins.

| Software and plugin | Source |
| --- | --- |
| Python | https://www.python.org/ |
| Micro-Manager 2.0.1 | https://micro-manager.org/ |
| FIJI distribution of ImageJ | https://imagej.net/software/ |
| TrackMate Plugin | https://imagej.net/plugins/trackmate/ |
| Graphpad Prism | [https:// https://www.graphpad.com/](https://www.graphpad.com/) |

## Movie Legends

**Movie 1: aPKC and ZO-1 colocalize at the apical cell-cell junction**

Z series through a polarized mESC spheroid with endogenously tagged mNG::aPKCι (green) and ZO-1::HaloTag labeled with JF646 (magenta). Z slices are separated by 1 µm.

**Movie 2: ZO-1 clusters move apically during polarity establishment**

Time lapse of a polarizing mESC spheroid with endogenously tagged ZO-1::mNG with frames recorded at 15-second intervals.

**Movie 3: ZO-1 and F-actin punctae move together**

Time lapse of a polarizing mESC spheroid with endogenously tagged ZO-1::mNG (green) and labeled with SPY650-FastAct F-actin dye (magenta). Time stamp indicates minutes : seconds with frames recorded at 30-second intervals.

**Movie 4: ZO-1 retracts from the apical membrane upon MLCK inhibition**

Time lapse of an mESC spheroid with endogenously tagged ZO-1::mNG and treated with ML-7 at the start of the movie. Time stamp indicates minutes : seconds with frames recorded at 15-second intervals.

**Movie 5: Recovery of apical ZO-1 transport after ML-7 washout**

Time lapse of an mESC spheroid with endogenously tagged ZO-1::mNG after ML-7 containing medium was replaced with fresh medium at the start of the movie. Time stamp indicates minutes : seconds with frames recorded at 60-second intervals.

**Movie 6: Apical PCX localization is disrupted following MLCK inhibition**

Time lapse of an mESC spheroid expressing transgenic PCX::mNG and treated with ML-7 at the start of the movie. Time stamp indicates minutes : seconds with frames recorded at 15-second intervals.

**Movie 7: F-actin is redistributed following MLCK inhibition**

Time lapse of an mESC spheroid with endogenously tagged ZO-1::mNG (not shown) and labeled with SPY650-FastAct F-actin dye, treated with ML-7 at the start of the movie. Time stamp indicates minutes : seconds with frames recorded at 60-second intervals.

**Movie 8: Dual effects of H1152 on ZO-1 localization on different timescales**

Time lapse of an mESC spheroid with endogenously tagged ZO-1::mNG and treated with H1152 at the start of the movie. Note that ZO-1 particles continue moving apically after H1152 treatment, but after ∼35 minutes of exposure to the drug, an ectopic pool begins to accumulate basally (arrowhead). Time stamp indicates minutes : seconds with frames recorded at 60-second intervals.

